# Spatially-resolved multimodal profiling identifies functionally-heterogeneous cancer-associated fibroblasts associated with poor radiotherapy outcomes in muscle-invasive bladder cancer

**DOI:** 10.64898/2026.09.18.752662

**Authors:** Amy Burley, Katie O’Fee, Morgane Dromby, Mi Chween Chan, Barry Gusterson, Adrian Larkeryd, Emma Westlund, Priya Narayanan, Chirine Sakr, Tatiany Silveira, Rose Foster, Mikel Portillo, Marc Rabionet Diaz, Tom Lund, Alan Melcher, Esther N. Arwert, Richard Bryan, Maurice Zeegers, K.K. Cheng, Ananya Choudhury, Kelly Jones, Shaista Hafeez, Robert Huddart, Syed Hussain, Emma Hall, Nick James, Anna Wilkins

## Abstract

**Background:** Cancer-associated fibroblasts (CAFs) contribute to systemic therapy resistance in muscle-invasive bladder cancer (MIBC), but their functional heterogeneity and relevance to curative bladder-preserving radiotherapy, are poorly understood.

**Methods:** Transcriptomic analysis was performed on 279 tumours from BC2001, a phase 3 radiotherapy clinical trial. To study CAF heterogeneity, we integrated bulk RNA-seq, single-cell spatial analysis of multiplex immunofluorescence images and quantification of extracellular matrix (ECM) features in 155 MIBC biopsies. The functional heterogeneity of distinct CAF populations was evaluated by single-nuclear RNA-seq. Spatial interactions between CAF populations and CD8+ T-cells was assessed and the relevance of lymphocytes to radiation responses was evaluated in a CAF-enriched murine bladder cancer model (BBN963).

**Results:** BC2001 patients with CAF-enriched tumours had worse overall survival (HR=1.671, 95% CI 1.221-2.287, Log-rank p=0.0012). CAF abundance and antigen expression was highly heterogenous. Podoplanin (PDPN) was expressed on the majority of CAFs and was associated with inflammatory pathways. Enrichment of CAF gene signatures was associated with a significant increase in CAFs expressing fibroblast activation protein (FAP) (p=0.004) and dense ECM features (p=0.0004). Fifty-three percent of tumours exhibited stromal CD8+ T-cell exclusion with significant enrichment in FAP-dominant neighbourhoods (p<0.001). *In vivo*, lymphocytes were critical for radiation-induced tumour control, indicating that immune cold or excluded tumours may have limited radiotherapy responses.

**Conclusion:** In MIBC, CAFs are associated with poor radiotherapy outcomes. Multiple mechanisms are deployed by functionally-heterogeneous CAFs, including promotion of chronic inflammation by PDPN+ CAFs and ECM remodelling by FAP+ CAFs which impact CD8+ T-cell distribution and radiation responses.

**Graphical abstract:** 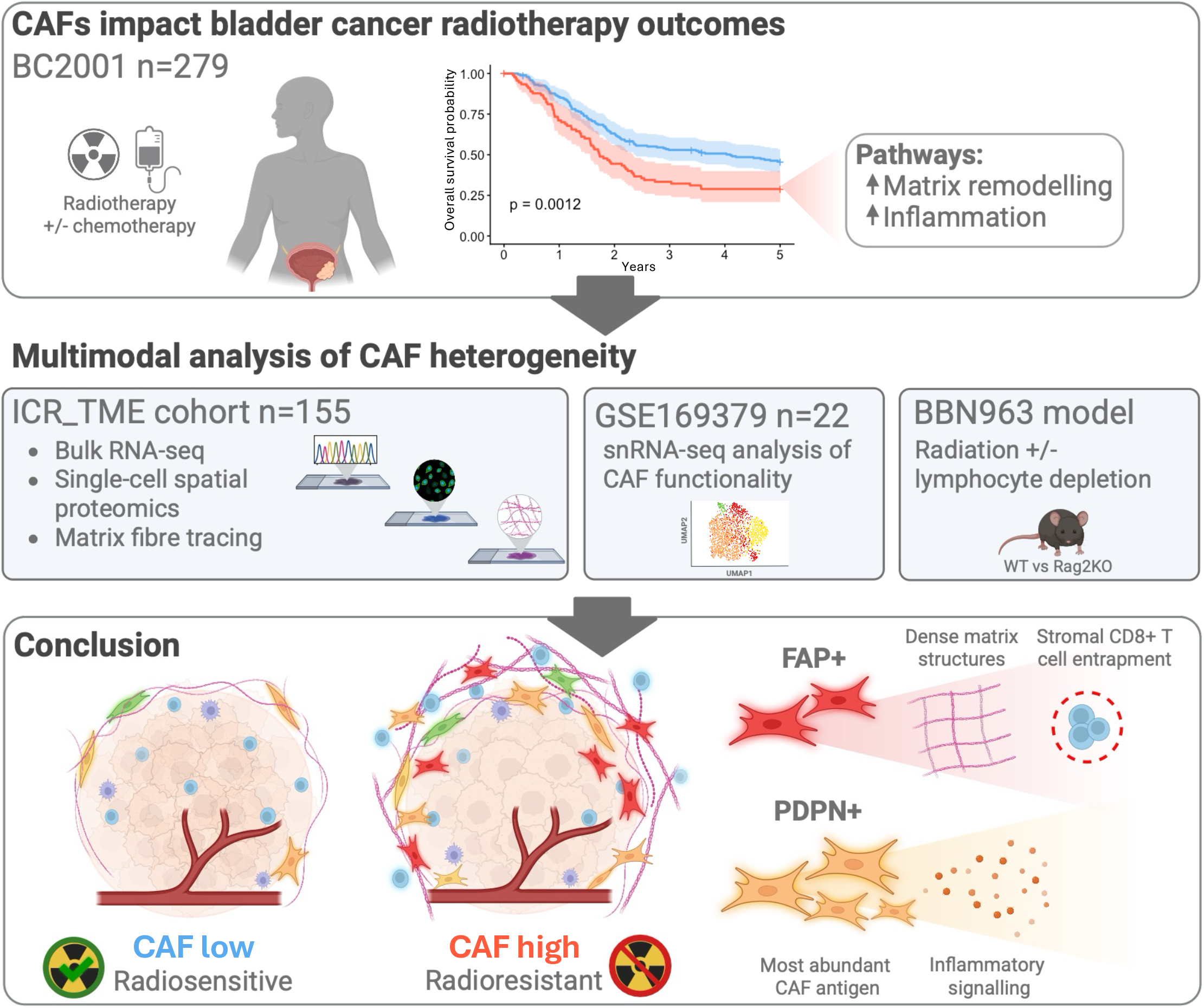

## Introduction

Bladder cancer is the 8^th^ most common cancer worldwide with over 635,000 cases annually [1]. Muscle-invasive bladder cancer (MIBC) has an aggressive disease trajectory, for which five-year survival following curative-intent treatment has historically remained at ∼60% for patients with disease confined to the pelvis [2–4]. Novel systemic therapies, including the antibody drug conjugate/PD-1 inhibitor combination enfortumab vedotin/pembrolizumab have markedly improved survival outcomes and increased pathological complete response rates [5, 6]. As a result, there is a heightened interest in bladder preserving approaches whereby patients can avoid radical cystectomy and the associated quality-of-life detriment. Chemoradiotherapy is an established curative-intent, standard-of-care treatment for MIBC which enables bladder preservation [7], with advances in precision radiotherapy and the addition of novel systemic therapy further improving the therapeutic index of chemoradiation [8–10]. Despite these advances, treatment outcomes remain variable, highlighting the need to better understand the biological factors that influence response to chemoradiotherapy.

The tumour microenvironment (TME) comprises multiple non-malignant cell types including components of the adaptive and innate immune system, vasculature and structural elements such as cancer-associated fibroblasts (CAFs). The composition of the TME and interactions between cell types within it has the potential to influence treatment responses by a variety of mechanisms. The role of CAFs in the TME is complex, with many studies reporting diverse pro-tumoral functions such as remodelling of the extracellular matrix, recruitment of immunosuppressive immune cells and direct effects on tumour cell proliferation, stemness and migration [11]. CAFs do not represent a single functional cell population. Rather, they comprise a heterogenous population of stromal cells with a degree of plasticity whose state is shaped by the cell of origin, biomechanics of the local extracellular matrix (ECM), ligand gradients, and spatial cues from surrounding cells.

Early descriptions of CAF heterogeneity defined two broad CAF phenotypes: myofibroblastic CAFs (myCAFs), characterised by transforming growth factor beta (TGF-β) activation and high expression of alpha-smooth muscle actin (αSMA), and inflammatory CAFs (iCAFs), activated by interleukin-1 alpha (IL-1α) [13,14]. Further research has refined understanding of TGF-β-driven CAF activation as being specifically relevant to CAFs expressing Leucine-Rich Repeat Containing 15 (LRRC15) [12]. With advances in single-cell technologies, our knowledge of CAF functional heterogeneity has rapidly expanded beyond definitions of myCAFs and iCAFs to include additional phenotypes such as antigen-presenting CAFs, amongst others [11, 13]. In addition, the importance of the neighbouring cell populations that shape the spatial context in which CAFs reside is increasingly recognised [14]. In support of this, CAF-centred spatial organisational patterns or “archetypes” appear to be reasonably conserved across different cancer types [15].

In bladder cancer, CAFs have been associated with poor responses to chemotherapy and worse survival following surgery [16–19]. Moreover, CAF activation via TGF-β has been associated with poor immunotherapy response in patients with metastatic bladder cancer [20, 21]. At a mechanistic level, Mariathasan et al., demonstrated that CAF-derived peri-tumoral collagen sequesters effector CD8+ T-cells at the tumour periphery creating an immune-excluded, treatment-resistant TME [20]. Despite these insights, the prognostic relevance of CAFs in patients with MIBC treated with radiotherapy remains poorly defined. Moreover, a spatially-resolved understanding of CAF functional heterogeneity and their impact on CD8+ T-cell infiltration that may underpin treatment responses in bladder cancer is lacking.

Studies in other tumour types, including pancreatic and colorectal cancer, have shown direct links between CAFs and radioresistance. Relevant mechanisms include radiation-induced changes in the secretory profile of CAFs (such as an increase in the expression of inducible nitric oxide synthase (iNOS) and secretion of nitric oxide [22]), adoption of a senescence-like phenotype [23] and enhanced deposition of ECM structures [24]. Together, these mechanisms result in immunomodulation, immune exclusion and activation of signalling pathways that promote tumour cell growth and survival following irradiation.

These findings indicate a potential for therapeutic targeting of CAFs, including to enhance the efficacy of existing treatments such as radiotherapy. Despite some preclinical success, approaches targeting CAFs via the TGF-β axis have shown limited clinical efficacy [25–28], and targeting αSMA+ myCAFs resulted in accelerated tumour growth in some pre-clinical studies [29, 30]. Such findings highlight the context-dependent and potentially opposing functions of different CAF populations, and caution against considering CAFs as a single therapeutic entity. Whilst the subsequent expansion in novel CAF targeting approaches such as fibroblast activation protein (FAP)-directed IL2 variant Simlukafusp alpha [31, 32], inhibition of the CAF secretory proteins CXCL2 and IL-1 [33, 34], and CAF reprogramming [35–37] presents further opportunity to target a variety of CAF populations, previous negative results highlight the complexity of CAF heterogeneity and the need for greater biological understanding of how diverse CAF populations may influence therapy response [38–40]. In this context, we define the inter-tumoral and functional heterogeneity of CAFs in MIBC with spatial resolution and examine their association with clinical outcomes in the context of radiotherapy.

## Results

### CAF enrichment is associated with poor overall survival in a radiotherapy-treated MIBC cohort

To evaluate whether CAFs are relevant to survival after radiotherapy in MIBC, we performed transcriptomic analysis of 279 diagnostic tumour samples from patients recruited to BC2001, a practice-changing phase 3 trial of radiation/chemo-radiation [41, 42] (Figure 1A). We observed significantly worse overall survival in patients whose tumours had a high ESTIMATE Stromal score, a well-established signature of CAF abundance (Figure 1B and S1A) [43], which remained significant after adjustment for known clinical prognostic factors in MIBC (hazard ratio (HR) 1.73, 95% confidence interval (CI) 1.25 – 2.38, Log-rank p<0.001) (Figure 1C). The ESTIMATE Immune score was not associated with overall survival (Log-rank p=0.73) (Figure S1B), although the combination of ESTIMATE Stromal and Immune signatures enabled identification of a subset of patients with a “Stromal low Immune high” profile who had particularly good survival after radiation (Figure S1C). We confirmed these findings using additional cell type deconvolution [44] where we observed a highly significant association between the CAF signature and poor survival after radiation (HR=1.52, CI=1.12-2.06, Log-rank p=0.0073) (Figure S1D), but no association with signatures of CD8+ T-cells (Log-rank p=0.25) or monocyte/macrophage populations (Log-rank p=0.85) (Figure S1E-F).

**Figure 1:**
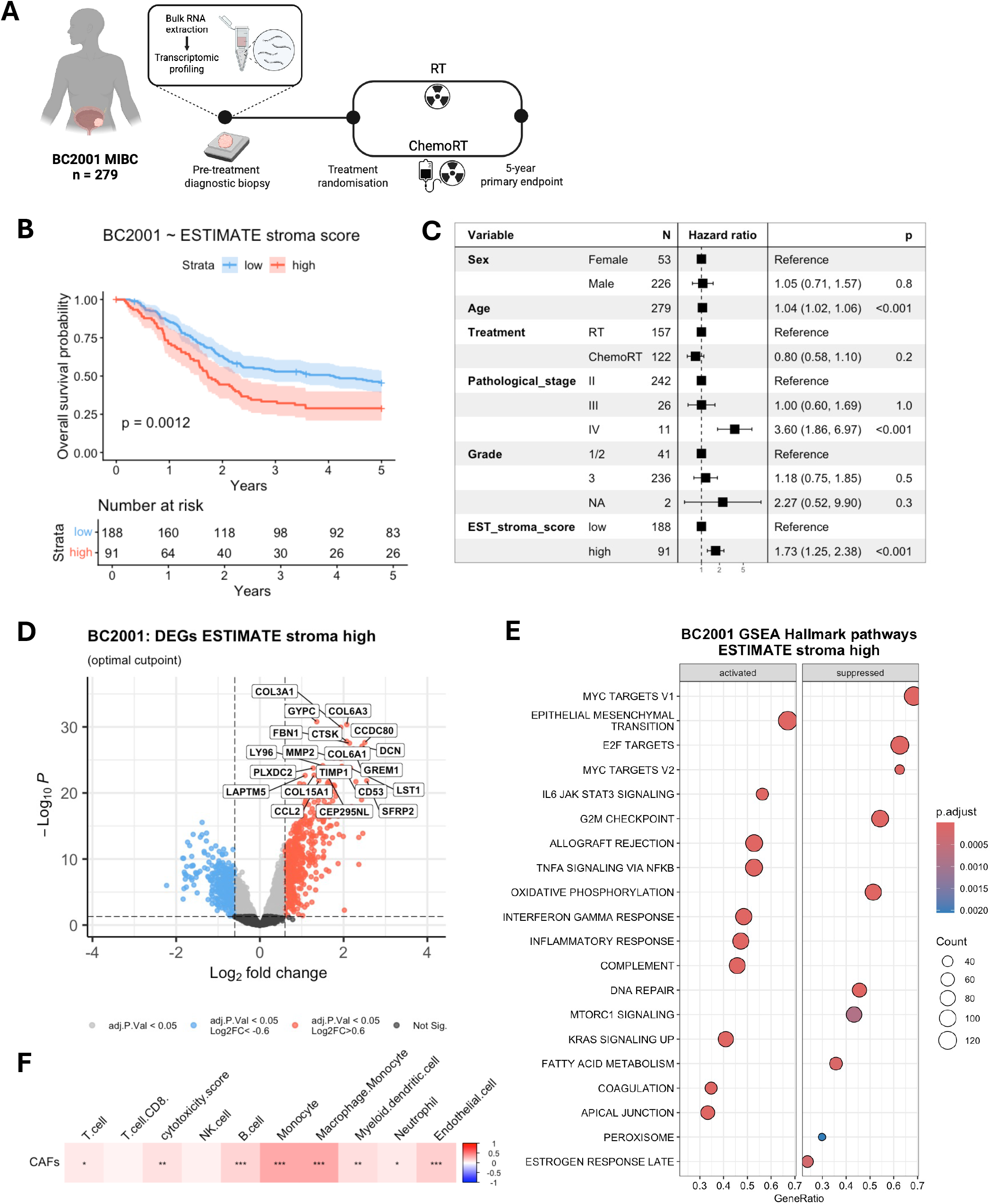
Stromal enrichment is associated with poor overall survival in BC2001, a radiotherapy treated MIBC trial cohort. **A**. A schematic of BC2001 sample collection and trial design. **B**. A Kaplan-Meier of BC2001 overall survival dependent on the ESTIMATE Stromal score at the optimal cut point, Stromal high hazard ratio = 1.671, 95% confidence intervals 1.221-2.287, log-rank p = 0.0012. **C**. A forest plot of the Cox multivariate analysis of overall survival. **D**. A volcano plot showing the top 20 differentially expressed genes upregulated in the Stromal high group (optimal cut point). **E**. A dot plot of the top hallmark pathways activated or suppressed in Stromal high samples (optimal cut point). **F**. The correlations between CAFs and other tumour microenvironment cell types as defined by Microenvironment Cell Populations (MCP)-counter cell deconvolution, (p<0.001 ***, p<0.01 **, p<0.05 *).

In order to better understand the biology underlying survival differences after radiation, according to CAF abundance, we performed differential gene expression analysis in ESTIMATE Stromal high (n=91) versus low (n=188) tumours. We identified that the top genes upregulated in Stromal high tumours included a number of genes related to collagens and the extracellular matrix remodelling (*MMP2*, *TIMP1*, *FBN1*) (Figure 1D). Subsequent gene set enrichment analysis (GSEA) identified that a number of inflammatory pathways were also significantly upregulated in ESTIMATE high tumours including IL6-JAK-STAT3 signalling, allograft rejection, interferon gamma response, TNFα signalling via NFκB and complement (Figure 1E and S1G).

We observed a strong positive correlation (R=0.57, p= <0.001) between ESTIMATE Stromal and Immune signatures suggesting that there were biological associations between CAF abundance and immune populations (Figure S1H). CAF-enriched tumours had significantly increased signatures for macrophage/monocytes and endothelial cells, as well as a smaller but significant correlation between CAFs and T-cells, B cells, neutrophils, and the cytotoxicity score (Figure 1F). As the ESTIMATE Immune, CD8 T-cell and macrophage signatures did not predict survival after radiation, this suggests that additional context regarding the location and status of the immune infiltrate is required and may relate to CAF abundance, location and heterogeneity. In summary, these findings indicate that CAF-enriched MIBC tumours have significantly worse responses to radiation and that biological features relating to both the extracellular matrix and inflammatory response are relevant to this inferior response.

### CAFs in MIBC are highly heterogenous with PDPN+ CAFs as the most common population

To build on the above findings and assess CAFs in the TME of MIBC more comprehensively, we performed multimodal profiling of an additional MIBC cohort containing baseline biopsies from 155 patients (the ICR_TME cohort). To evaluate heterogenous CAF populations and their spatial relationship with tumour and CD8+ T-cells, we applied a 6-marker multiplex immunofluorescence (mIF) panel that incorporated 4 frequently cited CAF antigens associated with functionally distinct mechanisms including: myCAF marker, alpha smooth muscle actin (αSMA) [45], marker of fibroblast activation, fibroblast activation protein (FAP) [46] and iCAF markers podoplanin (PDPN) [47] and platelet-derived growth factor receptor alpha (PDGFRα) [48, 49].

Qualitative assessment of multiplex images highlighted heterogenous morphological features and staining patterns across the ICR_TME cohort (Figure 2A top panel). Delineation of tumour, stroma and muscle regions enabled quantification of stromal abundance, and helped to distinguish αSMA+ CAFs in the stroma from muscle bundles and vasculature (Figure 2A middle panel). Stromal abundance was highly variable (range = 4.77 - 70.25%, mean 29.09%) (Figure 2B and S2C). Samples with a high ESTIMATE Stromal score had a highly significant enrichment in the percentage of stromal pixels detected in the multiplex images (Wilcoxon p=1.8e-08), confirming concordance between transcriptomic and proteomic approaches (Figure 2C).

**Figure 2:**
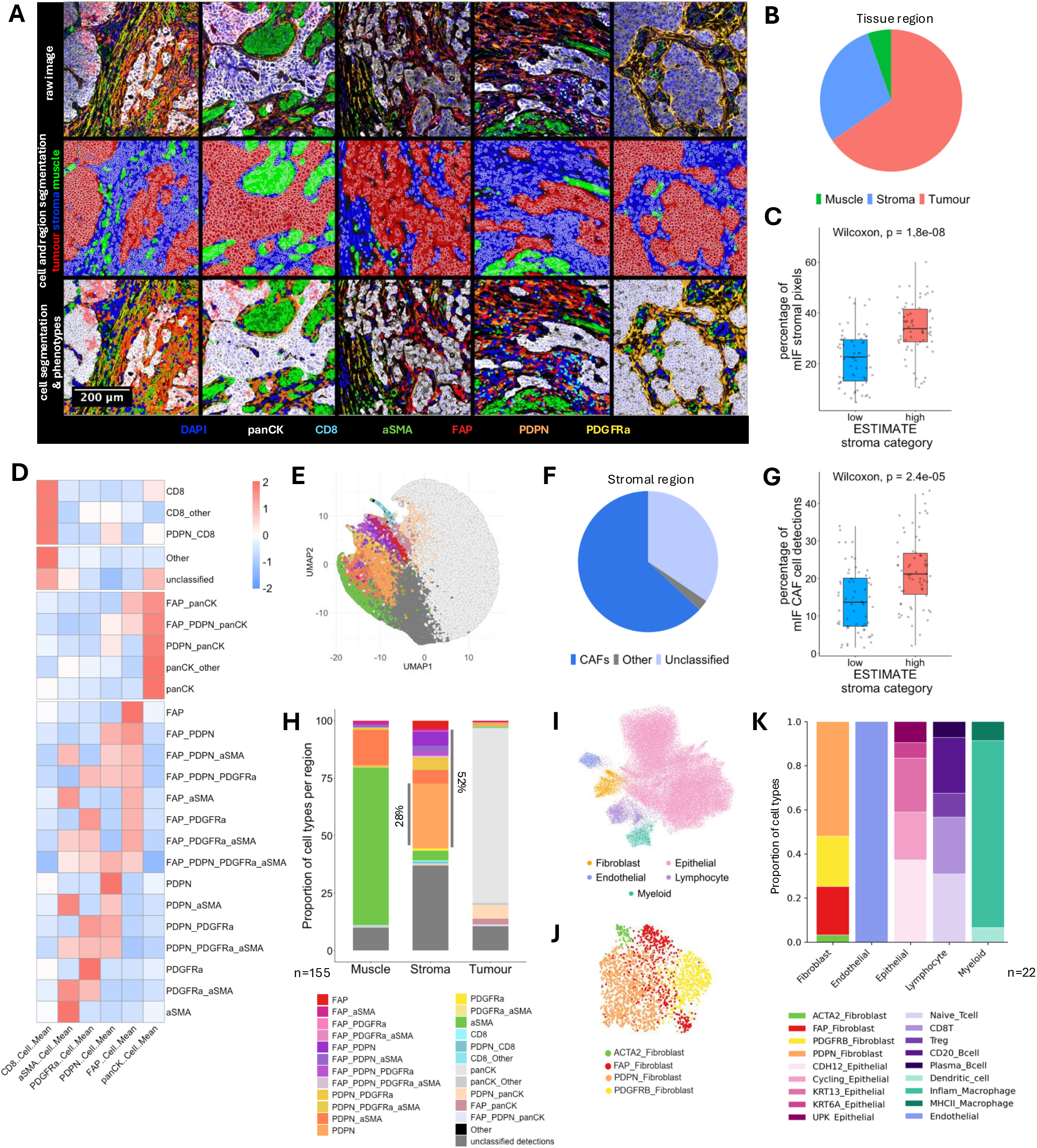
Single-cell analysis of multiplex immunofluorescence images highlights the abundance of distinct CAF populations in MIBC. **A. Top panel:** Five independent MIBC samples stained with the CAF 6-plex immunofluorescence panel, highlighting the expression of pan-cytokeratin (panCK), CD8, alpha Smooth Muscle Actin (αSMA), Fibroblast Activation Protein (FAP), Podoplanin (PDPN) and Platelet-Derived Growth Factor Receptor alpha (PDGFRα). **Middle panel:** Examples of Stardist cell segmentation plus the QuPath pixel classifier showing regional demarcation of tumour (red), stroma (blue) and muscle bundles and vasculature (green). **Bottom panel:** Segmented cells, coloured by cell phenotype as assigned by the QuPath composite object classifier. **B**. The mean percentage of pixels classified as tumour, stroma or muscle across all images in the cohort n = 155. **C**. A boxplot of the percentage of stromal pixels identified in the multiplex images of samples defined as ESTIMATE Stromal low or high (n=124) (Wilcoxon p=1.8e-08). **D**. A heatmap of the assigned cell phenotypes and their scaled cell mean expression of each antigen on the 6-plex panel. **E**. A UMAP of 100,000 randomly selected cells clustered by cell mean intensity with the assigned cell phenotypes displayed. Phenotype key as per 2H. **F**. The mean percentage of stromal cells classified as CAFs, unclassified stromal cells or other (i.e. CD8+ T-cells). **G**. A boxplot of the percentage of cells classified as CAFs in the multiplex images of samples defined as ESTIMATE Stromal low or high (n=124) (Wilcoxon p=2.4e-05). **H**. Quantification of the proportion of each cell type present in regions of tumour, stroma and muscle. **I**. UMAP embedding of GSE169379 single-nuclear RNA sequencing data, clustered by gene expression data with assigned cell phenotypes displayed. **J**. UMAP embedding of fibroblast populations in GSE169379. **K.** Quantification of the proportion of cell subtypes present in GSE169379.

Next, using cell segmentation and phenotypic assignment in QuPath (see methods) we assessed the cell type composition and spatial arrangement of >73 million cells at single cell resolution (Figure 2A bottom panel and S2B). We identified 25 cell phenotypes including tumour cells, CD8+ T-cells and 15 CAF populations expressing various combinations of αSMA, FAP, PDPN and PDGFRα. QuPath-assigned phenotypes showed expected concordance with antigen intensity (Figure 2D). Dimensionality reduction clustered cells according to phenotype and showed the relative proportion of cell types present in the cohort (Figure 2E and S2E).

Considering all CAF populations identified, we observed that CAFs were a dominant feature of the stroma in MIBC. On average, 63% of stromal cells were classified as CAFs (range 2.67 – 93.36%) (Figure 2F and S2D). Using a further mIF TME panel and a 66-plex PhenoCycler panel on a subset samples, we confirmed that the remaining unclassified stromal cells represented a range of immune cells and vasculature not detected in the CAF mIF panel (Figure S3). The percentage of CAFs detected in the multiplex images was significantly enriched in samples assigned as ESTIMATE Stromal high (Wilcoxon p=2.4e-05), confirming that the ESTIMATE Stromal score is representative of CAF enrichment (Figure 2G).

Assessment of cell phenotypes by region revealed that PDPN+ CAFs were the most abundant CAF population in the stroma, with PDPN expressed alone in 28% of stromal cells, or cumulatively with PDPN expressed in combination with other CAF markers in 52% of stromal cells (Figure 2H and S2F). In contrast, αSMA+ CAFs, traditionally thought of as a classical marker of TGF-β driven myCAFs, only represent a minority of stromal cells (15.5%), with similar proportions of FAP+ CAF populations.

To confirm these findings, we evaluated an independent, publicly-available MIBC single-nuclear RNA sequencing (snRNAseq) dataset comprising 22 MIBC tumours (GSE169379) [18]. This showed similarity in the proportion of CAF populations to the ICR_TME cohort where again PDPN+ fibroblasts were the most abundant fibroblast population, FAP+ CAFs comprised a substantial minority of CAFs, and αSMA+ (*ACTA2*) CAFs formed a small proportion (Figure 2 I-K and S2G). As PDPN is potentially expressed on different cell types in the stroma, we evaluated PDPN expression across major cell lineages in the TME in our 66-plex PhenoCycler panel. Consistent with our 6-plex evaluation, PDPN had the highest and most predominant expression in fibroblasts (Figure S3C-F).

### CAF-enriched MIBC tumours show specific enrichment of FAP+ CAFs

We next evaluated if specific patterns of inter-tumoral CAF heterogeneity were evident. Our results show that the differential abundance of distinct CAF populations was highly variable between each of the 155 MIBC tumours (Figure 3A). Some samples were predominantly composed of PDPN+ CAFs, whilst in other samples we observed that the majority of CAFs were FAP+ (predominantly FAP and FAP:PDPN CAF phenotypes).

**Figure 3:**
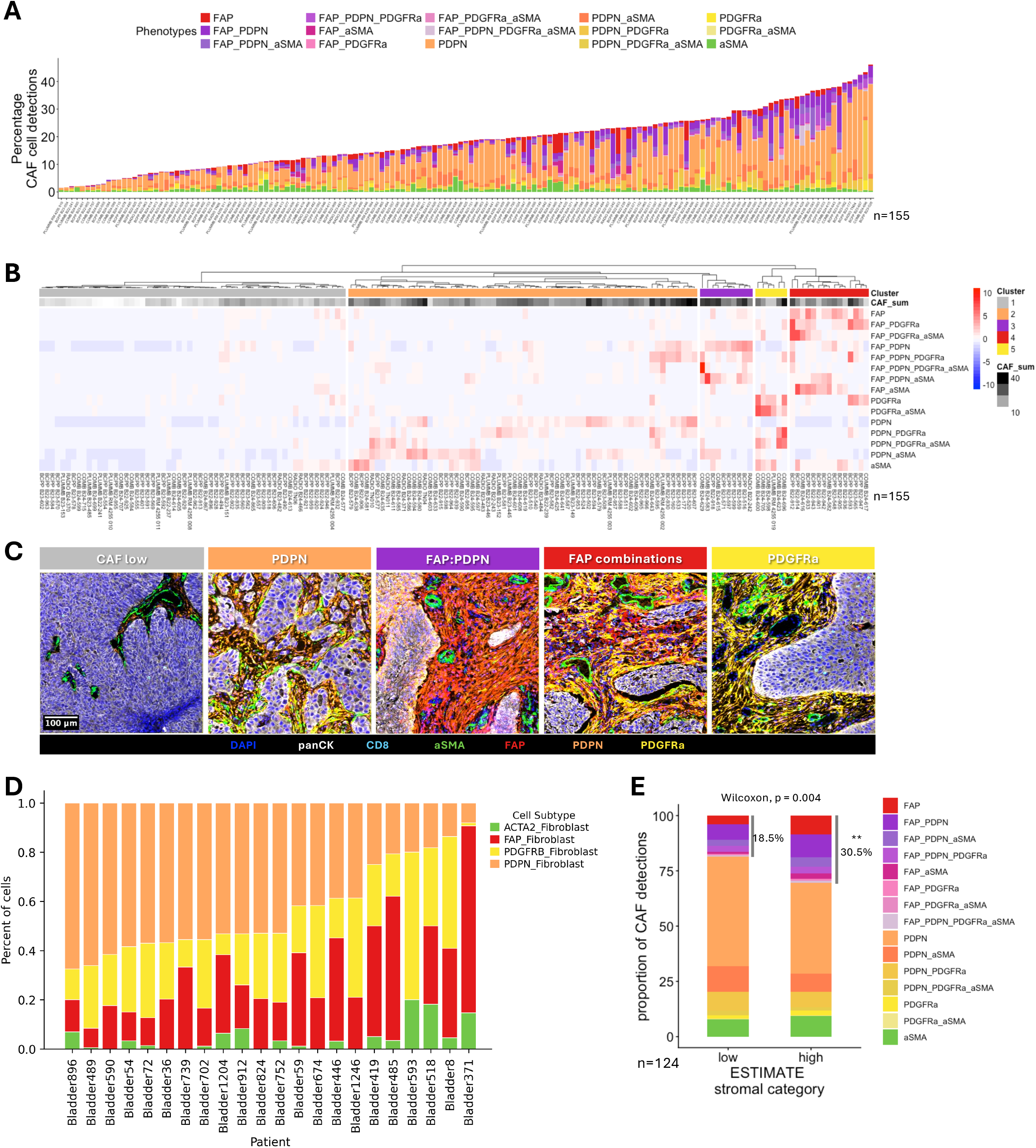
The composition of CAFs in the MIBC tumour microenvironment is highly heterogenous. **A**. The percentage of CAFs present in each sample, bars are filled with the relative proportion of CAF populations. **B**. A heatmap of samples clustered by CAF composition, annotated by the total percentage of CAFs detected in the multiplex image. **C.** Representative images of samples assigned to each CAF cluster. **D**. The proportion of fibroblast populations present in each sample in GSE169379. **E**. The mean proportion of CAF populations present in samples assigned as ESTIMATE Stromal low or high (n=124). Stars (**) indicate the significant difference in the proportion of combined FAP+ CAF subtypes as per supplementary figure 4C (Wilcoxon p = 0.004).

To evaluate this quantitatively, we performed unsupervised clustering of samples by CAF composition which identified 5 distinct sample clusters (Figure 3B). Cluster 1 represented samples with minimal CAF enrichment (37.4% of samples). The largest cluster (cluster 2, 42.6% of samples) was defined by an enrichment of PDPN+ CAFs expressed alone or in combination with αSMA. Clusters 3 and 4 were defined by an abundance of FAP+ CAF phenotypes, where cluster 3 represented samples with FAP:PDPN+ CAFs and cluster 4 was dominated by FAP+ CAF populations including FAP alone, FAP:PDGFRα+ and FAP:αSMA+ CAF phenotypes. Cluster 5 represented a small proportion of samples enriched for PDGFRα+ CAFs. Representative images of samples assigned to each cluster are shown in Figures 3C and S4A. Similar inter-tumoral diversity in the CAF composition was observed in the snRNAseq dataset with many tumours primarily composed of PDPN+ CAFs but a limited number containing predominantly FAP+ CAFs (Figure 3D).

Quantification of the proportion of CAF populations present in samples assigned as ESTIMATE Stromal low or high revealed a significant enrichment of FAP+ CAF populations in ESTIMATE Stromal high tumours (Wilcoxon p=0.004) (Figures 3E and S4B-C). In contrast, the relative proportion of multiple PDPN+ CAF populations decreased in ESTIMATE Stromal high tumours (Figure S4B).

### PDPN+ CAFs and FAP+ CAFs form spatially distinct neighbourhoods in the MIBC tumour microenvironment

To assess the spatial dynamics of the tumour microenvironment, we enlisted MuSpAn to identify recurrent cellular neighbourhoods (RCNs) across 143 samples stained with the CAF mIF panel (Figures 4A-B) [50]. We identified 8 RCNs, 3 of which represented tumour nests (RCN0, RCN3, RCN5), with RCN5 displaying an increase in the proportion of PDPN:panCK+ tumour cells. The remaining RCNs reflected stromal neighbourhoods. PDPN- and FAP-dominant CAFs formed almost exclusive spatial niches. RCN1 and RCN7 were dominated by PDPN+ CAFs, notably RCN1 represents PDPN+ CAFs interacting with unclassified cells. In agreement with our 66-plex PhenoCycler quantification (Figure S3), we speculate that these unclassified cells may reflect a combination of immune cells. RCN4 was dominated by a variety of FAP+ CAF phenotypes. Cross referencing RCN assignment with raw mIF images confirmed that RCN2 represented regions of muscle bundles and vasculature (Figure 4C and S5A). We did not observe fibroblast-like stromal RCNs with a distinct dominance of αSMA+ or PDGFRα+ CAFs.

**Figure 4:**
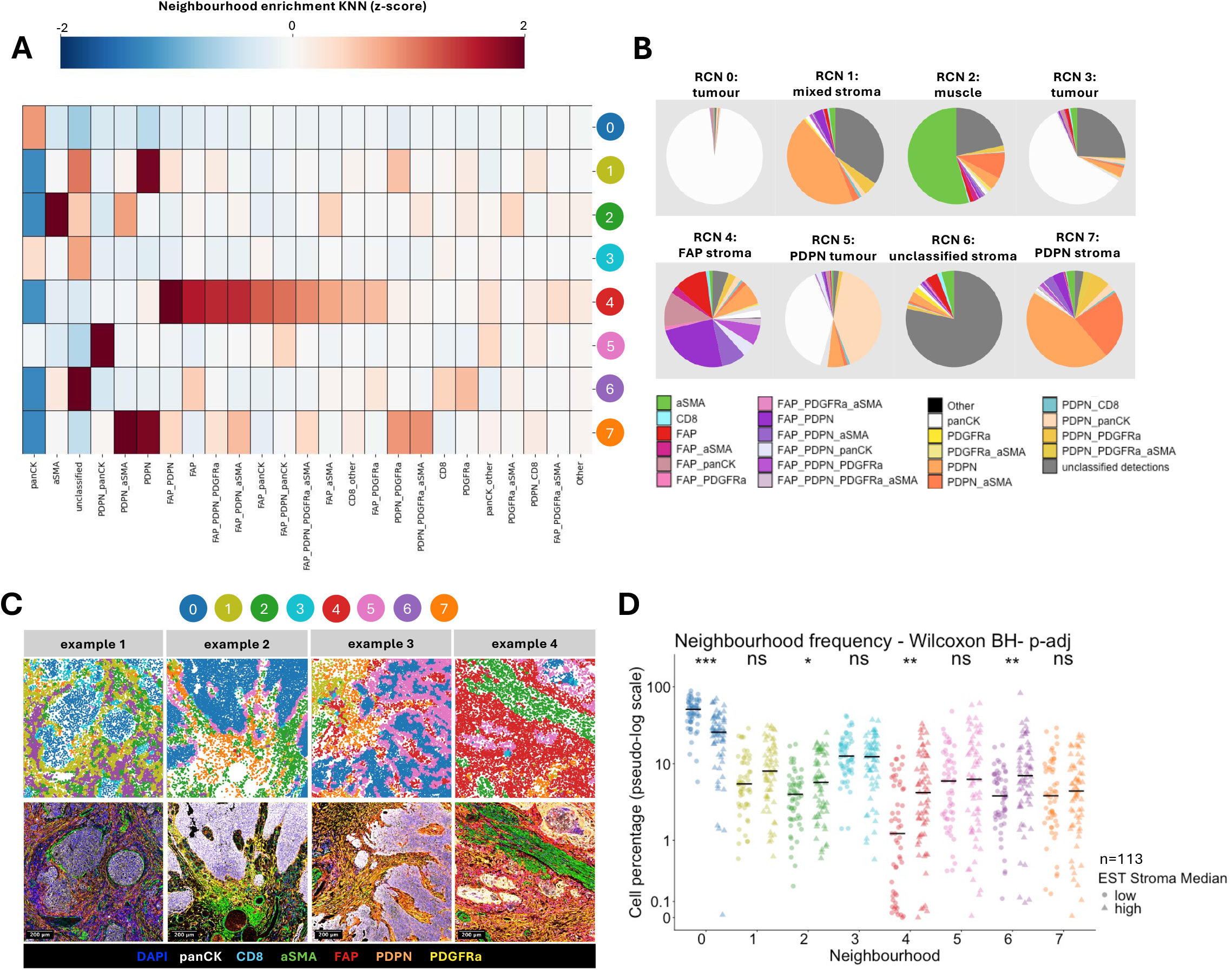
CAF populations form spatially distinct recurrent cellular neighbourhoods. **A.** A heatmap of the cell phenotypes that are enriched in the 8 recurrent cellular neighbourhoods (RCNs). **B**. Pie charts display the proportion of cell phenotypes assigned to each of the recurrent cellular neighbourhoods. **C**. Representative images of MIBC samples with cell centroids coloured by the assigned recurrent cellular neighbourhood (top panel) and the corresponding multiplex immunofluorescence image (bottom panel). **D**. A comparison of the neighbourhood frequency in samples assigned as ESTIMATE Stromal low or high, n=113 (low n=54, high n=59). Benjamini-Hochberg adjusted Wilcoxon p symbols are displayed, p<0.001 ***, p<0.01 **, p<0.05 *.

When assessing interactions between tumour and stromal RCNs, we observed that PDPN:panCK+ tumour cells assigned to RCN5 were often surrounded by FAP-dominant RCN4 as demonstrated in the Figure 4C example 4. We hypothesise that the tumour cell expression of PDPN represents tumour cells with an increased migratory profile that may in part be promoted by crosstalk with FAP+ CAFs [51–53].

For every sample, we assessed the percentage of cells assigned to each RCN and compared samples assigned as ESTIMATE Stromal low or high (Figure 4D). As expected, Stromal high samples had significantly less cells assigned to the dominant tumour nest RCN0 (Wilcoxon p<0.001). Consistent with our earlier observations, Stromal high samples had a significant enrichment in the percentage of cells assigned to the FAP-dominant RCN4 (Wilcoxon p=0.002), plus RCN2 (Wilcoxon p=0.04) and RCN6 (Wilcoxon p=0.004). These findings indicate that Stromal high samples are characterised by dense networks of FAP+ CAFs, plus increased muscle invasiveness and abundance of unclassified cells, which may represent immune populations, for example macrophages (Figure S3).

### PDPN+ CAFs and FAP+ CAFs show functional divergence in MIBC

Given the prominence of PDPN and FAP-dominant RCNs, we set out to determine if these CAF populations are functionally distinct. As these CAF populations were also present in the Gouin et al., snRNA seq dataset we assessed the top differentially-expressed genes that defined them (Figure 5A). FAP+ CAFs were specifically associated with expression of a number of genes relating to the ECM, including collagens (*COL1A1*, *COL1A2*), periostin (*POSTN*) and fibronectin (*FN1*). In contrast PDPN+ CAFs did not show specific expression of ECM genes. Instead, pathways analysis indicated that PDPN+ CAFs were specifically associated with upregulation of several inflammatory pathways such as TNFα signalling via NFkB, IL-6/JAK/STAT3 signalling and interferon alpha and gamma responses, plus TGFβ signalling (Figure 5B). We note that there was a trend towards downregulation of a number of these inflammatory pathways in FAP+ CAFs (Figure 5C).

**Figure 5:**
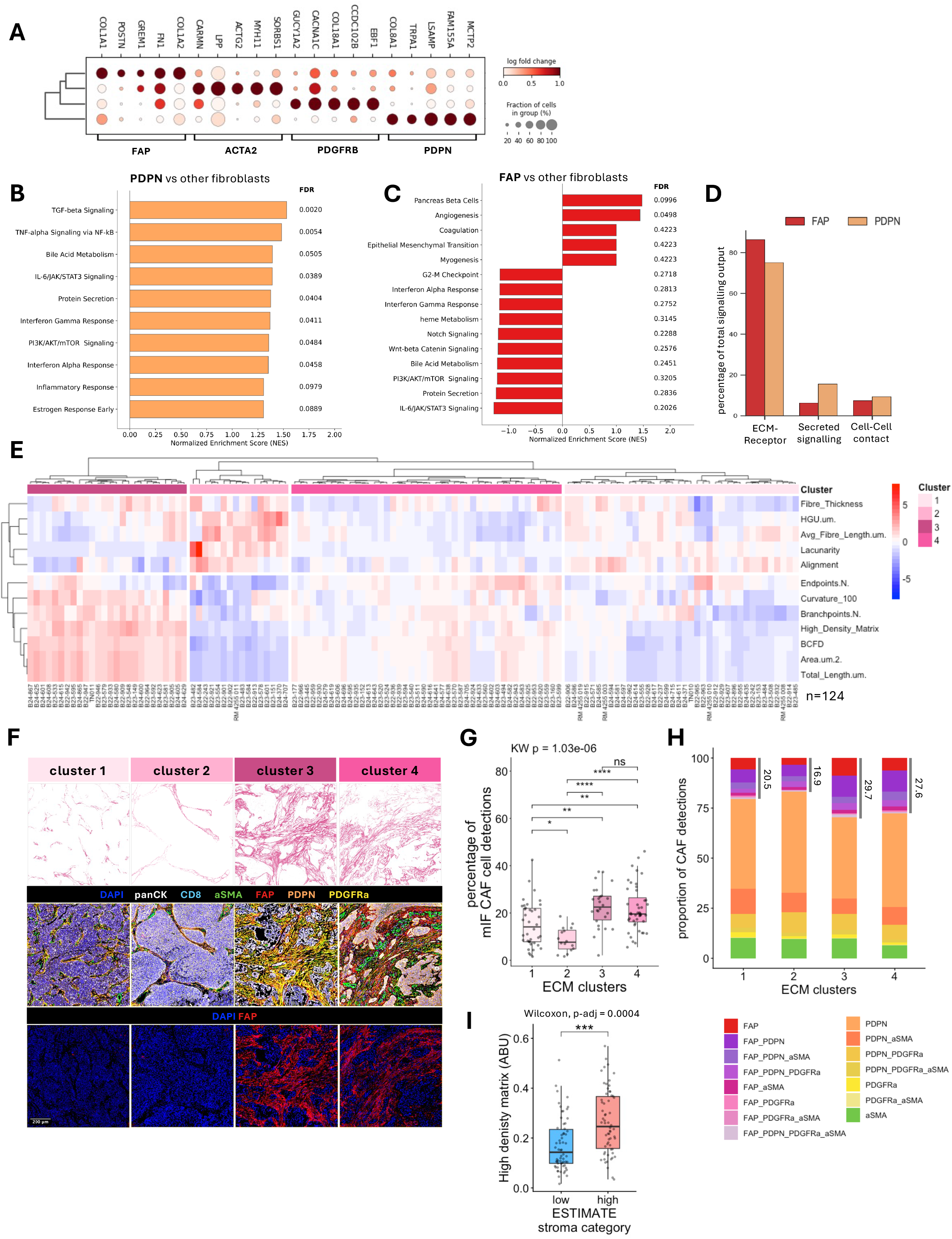
CAF populations have distinct functions. **A**. The top 5 differentially expressed genes associated with the four CAF populations identified in GSE169379. **B**. Normalised enrichment scores for top hallmark pathways that differentiate PDPN+ CAFs from all other fibroblasts in GSE169379. **C**. Normalised enrichment scores for top hallmark pathways that differentiate FAP+ CAFs from all other fibroblasts in GSE169379. **D**. A comparison of cell communication strategies (as defined by CellChat) in FAP and PDPN CAFs. **E**. A heatmap of ICR_TME samples clustered by extracellular matrix features quantified by TWOMBLI (n=124). **F**. Representative picrosirius red images of samples assigned to each ECM cluster (top panel), the corresponding multiplex image (middle panel) and FAP single-plex image (bottom panel). **G**. The percentage of CAFs present in samples assigned to each ECM cluster, Kruskal-Walis (KW) p-value = 1.03e-6, Benjamini-Hochberg adjusted Wilcoxon p symbols are displayed for pairwise comparisons, p<0.001 ***, p<0.01 **, p<0.05 *. **H**. The mean proportion of individual CAF populations present in samples assigned to each ECM cluster. **I**. A comparison of the high-density matrix score between samples assigned as ESTIMATE Stromal low or high, Benjamini-Hochberg adjusted Wilcoxon p = 0.0004, n=124.

We next assessed ligand receptor interactions of PDPN+ CAFs and FAP+ CAFs. As expected for cells of fibroblast lineage, both CAF populations largely communicated via interactions with the ECM, although this was more prevalent in FAP+ CAFs. In contrast, and consistent with the above pathways analysis, PDPN+ CAFs had an increased capacity to communicate via secreted signalling (Figure 5D).

To further assess functional divergence in FAP+ CAFs versus PDPN+ CAFs, we evaluated collagen ECM patterns in the ICR_TME cohort by picrosirius red staining of whole slide images and tile-based quantification of ECM features using the fibre tracing tool TWOMBLI (Figure S6A). 124 samples in the ICR_TME cohort had a complete set of mIF, ECM and transcriptomic data. Unsupervised clustering of ECM features revealed four distinct clusters of samples (Figure 5E). Clusters 1 and 2 have small quantities of collagen and can be further characterised by short and thin fibres (cluster 1) or vascular features (cluster 2). In contrast, clusters 3 and 4 have considerably more dense collagen deposition that forms heavily cross-linked ECM structures, this was particularly evident in cluster 3. Representative images of each cluster are shown in Figures 5F and S6B-C.

Combining ECM and mIF data confirmed that samples in ECM clusters 3 and 4 had significantly increased abundance of CAFs as quantified by mIF (Figure 5G). In addition, the formation of dense ECM structures in clusters 3 and 4 is associated with an enrichment of FAP+ CAFs and smaller proportion of PDPN+ CAFs (Figure 5H and S6D).

Combining ECM and transcriptomic data enabled us to evaluate ECM features according to ESTIMATE Stromal high versus low signatures. This confirmed that samples assigned as ESTIMATE Stromal high had significantly more high-density matrix (Wilcoxon p=0.0004), increased total fibre length (Wilcoxon p=0.0006), and increased complexity of ECM as indicated by higher fractal dimension (Wilcoxon p=0.0006) (Figure 5I and S6E). Overall, this suggests that ESTIMATE high samples have more dense and complex networks of collagen fibre structures, produced most commonly by FAP+ CAFs, contributing to a more desmoplastic and therapy-resistant TME.

### Spatial dynamics between CAFs and lymphocytes have important implications for radiation responses

With the inclusion of CD8 on the multiplex panel, we were able to assess the spatial distribution of CD8+ T-cells in the ICR_TME cohort and determine their interactions with the different CAF populations. First, we evaluated the proportion of CD8+ T-cells in stromal or tumour regions which revealed a highly significant enrichment of CD8+ T-cells in stromal regions compared to tumour nests (Wilcoxon test p= 2.2e-16) (Figure 6A). Using the CD8+ T-cell location, samples were stratified as CD8 desert, tumour nest excluded, tumour nest infiltrated or mixed. The tumour nest excluded category was the largest representing 53% of samples (Figure 6B-C and S7A). To further understand which cells may be responsible for CD8 sequestration in the stroma, we assessed CD8+ T-cell enrichment within our previously defined RCNs. Three stromal RCNs had a significant enrichment of CD8+ T-cells with the strongest enrichment found in the FAP-dominant RCN4 (p=9.5e-13) (Figure 6D). In addition, we assessed the frequency of nearest neighbour contacts between CD8+ T-cells (source) and different CAF populations or tumour cells (target). This revealed that CD8+ T-cells are frequently in contact with several FAP+ and PDGFRα+ CAF phenotypes. In contrast, CD8+ T-cells were less likely to be near CAFs that co-express αSMA (Figure S7B).

**Figure 6:**
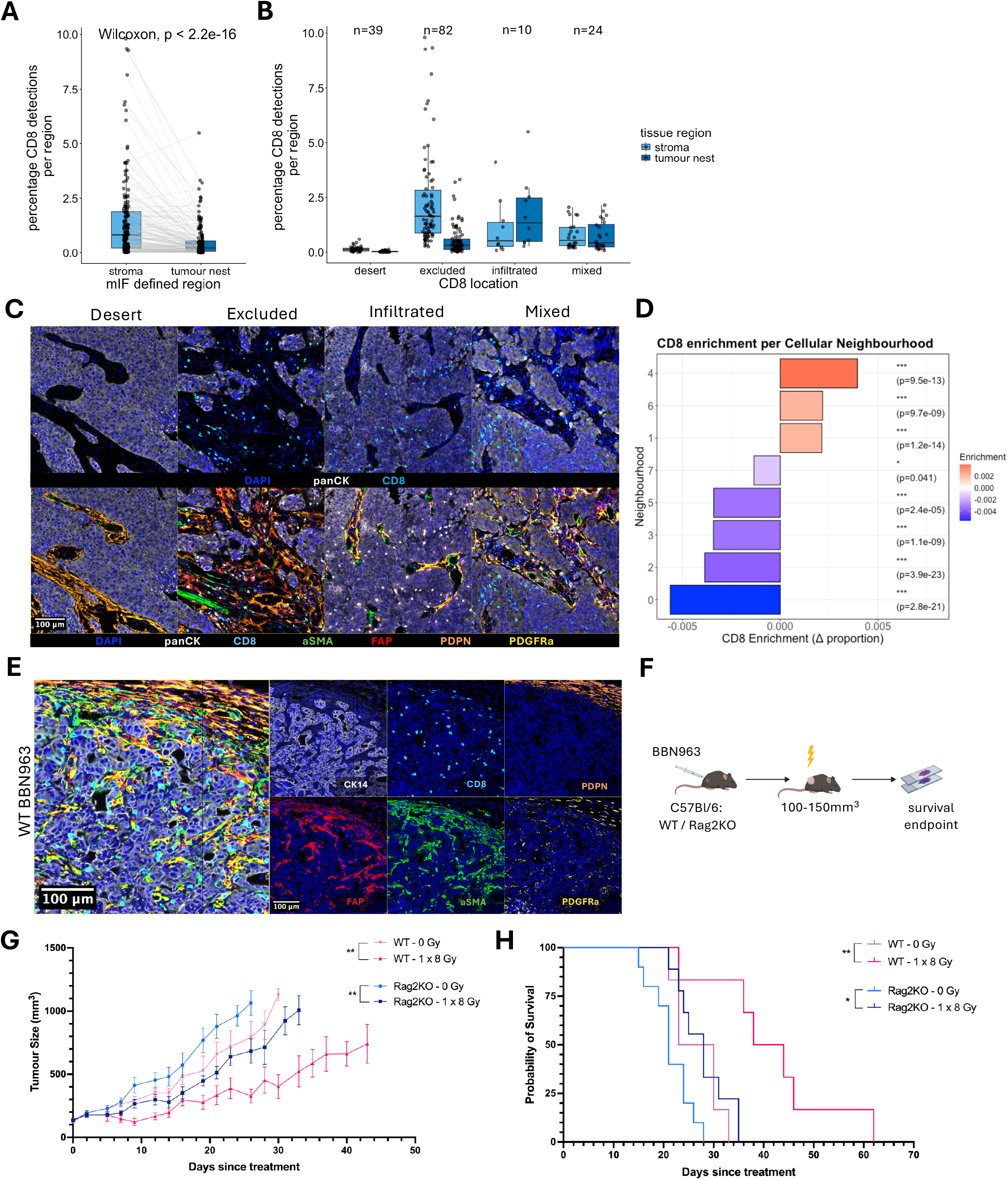
Spatial colocalisation of CAFs and CD8+ cells may impair radiation responses. **A**. A comparison of the percentage of CD8+ T-cells present in stroma or tumour nest regions in ICR_TME samples (as defined by the QuPath regional pixel classifier), Wilcoxon p = 2.2e-16. **B**. A comparison of the percentage of CD8+ T-cells present in stroma or tumour nest regions in samples defined as desert, tumour nest excluded, tumour nest infiltrated or mixed. **C**. Representative images of ICR_TME samples defined as desert, tumour nest excluded, tumour nest infiltrated and mixed. **D**. A comparison of CD8+ T-cell enrichment per recurrent cellular neighbourhood. Benjamini-Hochberg p adjusted values and symbols are displayed (p<0.001 ***, p<0.01 **, p<0.05 *) and indicate a significant enrichment of CD8+ cells inside a given RCN compared to outside the RCN (n=143). **E**. A multiplex immunofluorescence image of the murine-specific CAF panel in a wild-type (WT) BBN963 untreated tumour. **F**. A schematic of the BBN963 *in vivo* radiation study timeline. **G**. Tumour growth curves following 1 x 8 Gy radiation vs 0 Gy controls (WT 0 Gy vs 8 Gy, p=0.0051; Rag2KO 0 Gy vs 8 Gy, p=0.0065, Mixed-effects analysis). **H**. Survival curves following treatment (WT 0 Gy vs 8 Gy, p=0.0062; Rag2KO 0 Gy vs 8 Gy, p=0.0125, Log-rank (Mantel-Cox) test). Data was combined from 2 independent studies, (p<0.001 ***, p<0.01 **, p<0.05 *).

In metastatic bladder cancer, Mariathasan et al. showed that CD8+ T-cell exclusion in collagen-rich peri-tumoral regions was a feature of tumours with poor immunotherapy responses [21]. Taken together, our findings raise the possibility that patients with a cold or immune-excluded tumour microenvironment at baseline may have poor responses to radiotherapy. To assess the relevance of lymphocytes for the radiation response in MIBC, we established a syngeneic immunocompetent murine model of bladder cancer (BBN963). This model is known to recapitulate human basal-like bladder cancer in terms of responsiveness to immunotherapy with similar transcriptomic and histological features [54].

Using a murine-specific CAF multiplex immunofluorescence panel we confirmed that the BBN963 murine bladder cancer model had a stromal-rich tumour microenvironment with diverse CAF populations akin to the human disease (Figure 6E). In addition, the model showed fairly high CD8+ T-cell infiltration, as observed in human basal-like MIBC [54]. BBN963 cells were implanted subcutaneously into C57Bl/6 wild type (WT) mice or Rag2 knock-out (Rag2KO) mice which lack lymphocytes (Figure S7C). Once tumours reached 100-150 mm^3^ they were treated with 1 x 8 Gy using the small animal radiation research platform (SARRP) (Figure 6F and S7D). We observed a small proportion of spontaneous tumour regressions in the WT mice and slightly slower tumour growth in untreated WT versus Rag2KO mice (Figure S7E). With the addition of 1 x 8 Gy radiation, tumour growth was markedly slower with increased survival in WT mice versus Rag2KO mice (Figure 6G-H and S7F).

This confirms that lymphocytes are functionally relevant for radiation responses in MIBC. In view of the substantial CD8+ T-cell stromal sequestration, with particular enrichment in FAP+ CAF neighbourhoods, in combination with the functional importance of lymphocytes to radiation responses in MIBC, our findings indicate a potential therapeutic axis to improve radiation responses.

## Discussion

Understanding features of the tumour microenvironment that drive radioresistance is a clinical and scientific priority. We demonstrate for the first time that CAF enrichment is significantly associated with worse overall survival in a large, practice-changing phase 3 clinical trial of radiotherapy in MIBC. Previously, CAFs have been associated with an aggressive tumour trajectory and poor outcomes in non-muscle-invasive [55] and metastatic bladder cancer [21, 56], as well as following treatment with chemotherapy and surgery [16–19]. Our findings extend these observations by suggesting that heterogenous CAF populations also play an important role in radiotherapy response in MIBC and therefore provide a rationale for investigating CAF-targeting strategies as a means of improving the therapeutic index of radiation to enhance outcomes for patients with MIBC.

Given the diversity in CAF abundance that we observed, we suggest that CAF targeting approaches will be applicable to some patients, but not all. Moreover, the inter-patient heterogeneity in CAF populations, particularly with respect to PDPN+ and FAP+ CAFs, indicates that a one-size-fits-all CAF targeting approach is unlikely to be effective across all MIBCs. Intra-patient CAF heterogeneity is also an important consideration. Collectively, this may provide an explanation for why some CAF targeting approaches, including those acting on the TGFβ axis in MIBC, have previously failed [57]. We propose that some patients will be suitable for novel FAP-targeting approaches, whilst others may require approaches that manipulate the inflammatory signalling associated with PDPN+ CAFs, and some may require a combination. In addition, we acknowledge that CAF abundance and diversity is likely to change following radiation with impacts on tumour control and immune cell activity [47]. In this study, access to diagnostic samples provided insights into the TME at a single timepoint prior to any treatment. Comprehensive longitudinal profiling of CAFs in the context of the wider TME before, during and after radiation would be beneficial to further optimise CAF targeting with radiation.

Our deeper interrogation of the biology underpinning radioresistance in CAF-enriched MIBC highlighted roles of both the extracellular matrix remodelling and upregulation of several inflammatory pathways. Similar upregulation of interferon pathways, alongside IL6-JAK-STAT3, TNFα signalling via NFκB and other inflammatory pathways have been associated with poor radiotherapy outcomes in other tumour types [24, 58]. Collectively, these findings are consistent with chronic inflammation and interferon signalling in radiotherapy-refractory, fibroblast-enriched tumours, which we hypothesise is associated with T-cell dysfunction [59, 60]. A better understanding of these signalling pathways and associated heterotypic cellular interactions, including between CAFs and immune cells, could further inform rational development of novel targets alongside radiation in MIBC.

To understand the role of CAFs in MIBC, we evaluated 155 MIBC samples using a multimodal combination of transcriptomic analysis, single-cell spatial proteomic CAF profiling, and quantitative evaluation of the filamentous ECM. This is one of the largest studies to characterise CAFs in MIBC and contributes to greater spatially-resolved insights into CAF heterogeneity than what can be achieved via single marker immunohistochemistry (IHC) quantification or analysis of smaller single-cell RNA sequencing cohorts. Our results highlight a wide variation in CAF-enrichment across MIBC and identify PDPN+ CAFs as a particularly common population of CAFs with iCAF-like features. This is consistent with our previous work where we observed expansion of PDPN+ CAFs following radiation, in a process driven by inflammatory and interferon-related pathways and not by TGFβ signalling [47]. In healthy lymph nodes expression of PDPN on fibroblastic reticular cells is essential for immune regulation and stromal-immune cell crosstalk during the adaptive immune response [61, 62]. Taken together, we suggest that PDPN expression in CAFs plays a key role in immunomodulation of the TME. Further study of the inflammatory networks centred on PDPN+ CAFs, and downstream interactions with other immune cells in MIBC, would provide valuable insights for PDPN-directed CAF targeting approaches.

Spatial recurrent cellular neighbourhood analysis revealed that FAP+ and PDPN+ CAFs form almost exclusive stromal niches, as such we have focused our efforts on defining the functional properties of these major populations, while acknowledging that co-expression of multiple CAF antigens may be representative of phenotypic and functional plasticity. Our snRNA-seq analysis revealed that inflammatory pathways were not upregulated in FAP+ CAFs; instead, FAP+ CAFs were associated with genes relating to collagens and ECM remodelling. Consistent with these observations, tumour regions enriched for FAP+ CAFs were dominated by dense heavily crosslinked ECM structures which supports FAP+ CAFs functioning akin to myCAFs in MIBC. Tumours with dense ECM structures may be subject to greater mechanical forces resulting in further CAF activation via YAP [63], promotion of tumour cell growth, and activation of pro-tumoral Notch signalling [64, 65]. We note that patients with a high ESTIMATE Stromal score had significant enrichment of multiple FAP+ CAF populations and dense collagen ECM structures indicating that ECM-remodelling by FAP+ CAFs may be a key characteristic of radioresistant MIBC. Similar associations between FAP enrichment and treatment resistance were observed in the ABACUS trial of neo-adjuvant immunotherapy in MIBC [56]. In addition, FAP expression was associated with increased tumour stage and poor cancer-specific survival in high grade non-metastatic MIBC [66].

In the ICR_TME cohort, CD8+ T-cells showed prominent sequestration within stromal regions with minimal infiltration into tumour nests. This phenomenon could not be deciphered from bulk RNA-seq and highlights the importance of interrogating the spatial interactions between cells. Given the stromal sequestration of CD8+ T-cells in multiple CAF-enriched recurrent cellular neighbourhoods, particularly FAP-dominant RCN4, we propose that heterogenous CAF populations enlist multiple mechanisms to reduce CD8+ T-cell efficacy including physical restriction within dense ECM networks [67] and via immunosuppressive paracrine signalling resulting in reduced cytotoxicity and increased exhaustion [68, 69]. In view of the observed colocalisation of FAP+ CAFs and CD8+ T-cells in ICR_TME, and the emergence of novel FAP-directed CAF targeting approaches that incorporate T-cell stimulation [31, 33], we propose that FAP targeting could be an effective strategy to overcome CD8+ T-cell exclusion in patients with tumours enriched for FAP+ CAFs.

We acknowledge that CD8+ T-cell infiltration is likely to be impacted by multiple factors including tumour mutational burden and other immunomodulatory cell types in the TME. Likewise, CAFs are likely to impact the infiltration and activity of multiple lymphocyte populations, not just CD8+ T-cells. Genetic lymphocyte depletion in our *in vivo* bladder cancer murine model demonstrated the critical role of lymphocytes in tumour control following radiation. The significance of the immune infiltrate in radiation responses has been overlooked historically, and the synergistic effects of immunotherapy and radiotherapy remain unresolved. Clinical trials combining immunotherapy and radiotherapy have shown signs of early promise in MIBC [9, 10], but negative results have been reported in other tumour types [70, 71]. In view of our findings, we suggest that the quantity, location and spatial interactions of lymphocytes, in particular CD8+T-cells, should be an important consideration for future studies and the addition of CAF-targeting approaches has the potential to enhance synergistic effects.

In summary, we report a significant association between CAF enrichment and poor radiotherapy outcomes in MIBC. Multiple mechanisms are deployed by heterogeneous CAF subtypes in MIBC to promote radiation resistance which include promotion of chronic inflammation by PDPN+ CAFs and ECM remodelling by FAP+ CAFs. Rational selection of an optimal CAF target requires evaluation of the predominant CAF subtypes in each tumour. CAFs are a prominent feature of the MIBC tumour microenvironment and if this rational target selection can be accomplished, there is real potential for combined radiotherapy/CAF targeting to improve bladder-sparing therapeutic outcomes in MIBC.

## Methods

### Patient samples

BC2001 (ISRCTN68324339) was a randomised phase III study of radiotherapy with and without concomitant chemotherapy in muscle invasive bladder cancer [41, 42] (REC: 14/SC/1134). Samples were available for 279/458 trial participants. Clinical characteristics are provided in Table S1.

The ICR_TME cohort (n=155) is comprised of diagnostic transurethral resection of bladder tumour (TURBT) formalin-fixed paraffin-embedded (FFPE) samples collated from several clinical trials and tissue banks including: Bladder cancer prognosis programme (BCPP) [72] (REC: 06/MRE04/65), samples collected as part of the SELENIB trial (ISRCTN13889738); Collection of clinical material for molecular stratification in patients with muscle-invasive bladder cancer (COMB) (REC: 15/LO/0998); RAD-IO, a multi-stage randomised trial of durvalumab (Medi4736) with chemoradiotherapy with 5-fluorouracil and mitomycin C in patients with muscle-invasive bladder cancer [9] (ISRCTN43698103) and the Pembrolizumab in Muscle-invasive/Metastatic Bladder cancer Study (PLUMMB) (NCT02560636) which combines weekly radiotherapy with anti-PD-1 pembrolizumab for patients with advanced/metastatic bladder cancer [73] (REPEATS REC for RAD-IO and PLUMMB: 23/NE/0212).

### Sample processing

ICR_TME FFPE blocks were sectioned and stained with haematoxylin and eosin (H&E) to facilitate pathological review of material against the study inclusion criteria and to annotate regions of interest for analysis. Five 10 μm sections were cut for macrodissection and nucleic acid extraction from annotated regions, followed by 3 μm sections for multiplex imaging, and picrosirius red staining. A final H&E slide was stained to review the tissue remaining in the block (Figure S2A).

### Transcriptomic profiling

Transcriptomic data for BC2001 were generated, normalised and kindly shared with the ICR by Prof. Ananya Choudhury’s team at the University of Manchester. Briefly, data were acquired using RNA extracted from baseline tumour biopsies using an Affymetrix microarray to produce gene expression data for 20354 genes. The values were the GC-Content/Signal Space Transformation (GC/SST) normalised and the log2 expression of each gene in each sample was generated with Affymetrix’s apt-probeset-summarise and further normalised for plate batch effects by the function ComBat in the R package sva [74].

Transcriptomic profiling for BCPP samples was conducted by Veracyte. Data were normalised following the single channel array normalisation (SCAN) algorithm to adjust the GC content for each probe and to adjust the log2 raw intensity probe level data to subtract background noise.

RNA from COMB, RADIO and PLUMMB samples was sent to Genewiz (Azenta Life Sciences) for ribosomal RNA depletion, library preparation and strand-specific bulk RNA sequencing at 30M reads per sample. Fasta Quality (FASTQ) files were processed using the nf-core/rnaseq pipeline (https://zenodo.org/records/14537300). Alignment and quantification were conducted with Spliced Transcripts Alignment to a Reference (STAR) and RNA-Seq by Expectation Maximization (RSEM) respectively with reference to the human genome assembly GRCh38 Gencode Release 36 (GRCh38.p13).

### Bulk transcriptomic analysis

#### Cell deconvolution

ESTIMATE [43] and Microenvironment Cell Populations (MCP) cell deconvolution was conducted using the immunedeconv (v2.1.0) R package. Input data was a gene expression matrix of normalised, but not log-transformed count data, with rownames as HGNC gene symbols.

Correlations were calculated using the Kendall method to account for non-normally distributed data, p values were adjusted for multiple testing using the Benjamini-Hochberg method. Correlation heatmaps were generated using the corrplot R package (v0.95). The ESTIMATE Immune vs Stroma score scatter plot was generated with sm_statCorr from smplot2 (v0.2.5).

Due to differences in transcriptomic profiling, the ESTIMATE Stromal score and median cutpoint were calculated for each trial cohort within ICR_TME independently.

#### BC2001 Survival analysis

In BC2001 the key endpoint of interest for this study was overall survival (OS) defined as time from the date of randomisation to the date of death due to any cause. Patients alive at their last known follow-up were censored.

Survival analysis was conducted using the survfit function from the survival R package (v3.5-8). Kaplan-Meier curves were created using the ggsurvplot function from the survminer R package (v0.5.0). The surv_cutpoint function from the survminer package was used to find the optimal cutpoint for the ESTIMATE Stromal score in BC2001. The coxph function from the survival R package was used to fit cox proportional hazards regression models for multivariable analysis. Forest plots for Cox regression models were generated using the forestmodel R package (v0.6.2).

#### Differentially expressed genes and pathways analysis

A linear model was generated with Limma (v3.62.2) to compare samples assigned to groups of interest. Top differentially expressed genes were defined with an adjusted p value of 0.05 and Log2 fold change of 0.6. p values were adjusted for multiple testing using the Benjamini-Hochberg method. The volcano plot was produced with EnhancedVolcano (v1.24.0).

Gene set enrichment analysis was completed using the clusterProfiler R package (v4.14.6). Hallmarks (HM), Gene Ontology (GO) and Kyoto Encyclopedia of Genes and genomes (KEGG) (legacy) pathways were called using the msigdbr R package (v10.0.1). The Benjamini-Hochberg method was selected to adjust p values for multiple testing.

### PhenoImager image generation and quantification

#### Human multiplex staining

Prior to staining, slides were baked overnight at 50°C, followed by 1 hour at 60°C on the day of staining. All multiplex staining protocols were completed using the Leica BOND RX/RXm automated immunostainer and included the following steps: 1. tissue dewaxing and rehydration, 2. antigen retrieval, 3. H_2_0_2_ and protein blocking, 4. primary antibody staining, 5. secondary antibody staining, 6. application of Opal and all appropriate washes. Steps 2-6 were repeated for a further 5 cycles. The slides were manually washed and counterstained with DAPI before applying a coverslip. Antibody information for the human CAF and TME PhenoImager multiplex panels is provided in Table S2 and S3.

Multiplex immunofluorescence-stained slides were scanned on the PhenoImager (formerly Vectra Polaris v1.0.13) (Quanterix) at 20X. Following image acquisition, images were passed through the spectral unmixing software InForm (v2.6.0). Post InForm the output component image tiles were stitched to recreate the whole slide image using a stitching script generated by Pete Bankhead for QuPath (supplementary file 1).

#### Multiplex image analysis

Image visualisation and analysis was conducted in QuPath (v0.5.1) [75]. Regions of interest (ROIs) were generated following the pathologist-guided annotations conducted on H&E images. As per the cutting schema (Figure S2A), the sections for multiplex imaging were ∼50-60mm deeper into the block than the original annotated H&E section; therefore, annotations were adjusted to account for loss or gain of tissue. Staining and tissue artefacts incurred during the staining and imaging process were excluded from annotations. In cases where artefacts left insufficient tissue, the sample was excluded from further analysis. All annotations underwent a final pathologist review before entering the analysis pipeline.

A composite training image was generated from 52 randomly selected 1000 x 1000-pixel tiles from 32 independent whole-slide images. A pixel classifier was trained as an artificial neural network to detect and measure regions of tumour, stroma, and muscle/vasculature by providing examples of each region respectively. The classifier was also trained with an “ignore” category to reflect tissue areas that should not be classified, for example paucicellular areas or vessel lumens (supplementary file 2). Using the “Create Object” function the regions of tumour, stroma and muscle were made into annotations with the following conditions: minimum object size 150, minimum hole size 150.

Cell segmentation was achieved using the Stardist cell segmentation plugin for QuPath (Stardist v0.4.0)[76]. To phenotype the cellular detections, an object classifier was trained for each antigen, annotations were added to capture a wide variety of staining examples that reflect the heterogeneity of cells that were positive or negative for each antigen. Each classifier was generated using an artificial neural network. Single antigen object classifiers were compiled into a composite object classifier to facilitate detection of cell phenotypes that were positive for multiple antigens (supplementary file 3). In total, 64 cell phenotypes were identified. The phenotypes were refined to 25 based on known biology, observed phenotypes and abundance (supplementary file 4). A CAF was defined as a cell in the stromal region positive for any of the four CAF antigens and negative for pan-cytokeratin and CD8.

Three CD8 phenotypes were identified “CD8”, “PDPN:CD8” and “CD8 other” where PDPN:CD8 were an observed phenotype with coexpression of both antigens on small, round lymphocyte-like cells. “CD8 other” represent a small population of cells that co-express CD8 with αSMA, FAP or PDGFRα, these cells likely caused by limitations of cell segmentation in regions were cells are in close proximity.

To validate the refined phenotypes, a subset of 100,000 cells was randomly selected. For each phenotype the cell mean intensity for each marker was z-score normalised and the mean calculated. The scaled mean values were plotted against the assigned phenotype. Principal component analysis (PCA) was performed using the stats R package (v4.4.0) prcomp function. The first six principal components were retained and used as input for Uniform Manifold Approximation and Projection (UMAP). UMAP embeddings were generated using the uwot R package (v0.2.4) with the following parameters: n_neighbors = 10, min_dist = 0.05, spread = 2, n_components = 2 and metric = Euclidean.

The proportion of CAF phenotypes were compared in samples defined as ESTIMATE Stromal low (n=62) and stromal high (n=62). 15 CAF phenotypes were considered; Wilcoxon p values were calculated and adjusted for multiple testing with the Benjamini Hochberg method.

The percentage of CD8+ cells in tumour and stroma regions was defined as: (CD8+ cell count in the tumour region / total tumour region cell count) *100. (CD8+ cell count in the stroma region / total stromal region cell count) *100.

Samples were stratified into quartiles based on the total percentage of CD8+ T-cells. Samples in the lowest quartile were defined as “desert”. The remaining samples were classified as tumour nest excluded, tumour nest infiltrated or mixed based on the tumour: stroma CD8 ratio. Samples with a CD8 ratio > 0.2 were classified as “infiltrated”, those with a CD8 ratio < −0.2 were classified as “excluded”, and those with CD8 ratio values between −0.2 and 0.2 were classified as “mixed”.

### PhenoCycler image generation and quantification

Highly-plexed images were generated for 8 patient samples (2 samples per slide) with the PhenoCycler Fusion (v2.0, Quanterix) according to the manufacturers protocol, using the commercially available IO60 panel (Quanterix) plus 6 additional custom conjugations (Table S4). Cell segmentation was conducted in QuPath with the Stardist model (v.0.4.0). Cell-level data including antigen expression values, x and y coordinates, and sample meta data was exported.

Cells were filtered by size (50-200 µm^2^) to remove artefacts, >2.1 million cell were analysed. Antigen expression values were asinh normalised and scaled for each patient sample. Normalised data from each sample was combined into a single Spatial Experiment object using imcRtools (v1.16.0). Harmony data integration was performed by slide using selected antigens associated with phenotypic lineage (CD3e, CD11b, CD68, αSMA, CD31, Podoplanin, Vimentin, E-Cadherin and Pan-cytokeratin). To identify phenotypic clusters, Louvain community detection was conducted using Rphenograph (v0.99.) with k = 15. Clusters with similar antigen expression profiles were merged to define broad cell type metaclusters including epithelial, lymphocyte, myeloid, endothelial, muscle or fibroblasts. To refine the muscle metacluster in line with visual observations, the above analysis steps were repeated using the αSMA, Vimentin, Podoplanin, E-cadherin and Pan-Cytokeratin antigens.

Antigen expression values were visualised on the UMAP embedding using dittoDimPlot (v1.22.0). The lower and upper limits of the colour scale were determined by antigen expression values within the 1st and 99th percentiles to reduce the impact of outlier values. Podoplanin expression was considered across the assigned cell phenotypes. Cells with a normalised expression value >1 were considered as Podoplanin positive.

### Spatial analysis of multiplex images

The assigned cell phenotype and the cell centroid x and y coordinates were exported from n=143 PhenoImager images. To note, 12/155 ICR_TME samples were excluded from spatial analysis due to abundance of pan-cytokeratin negative tumour cells. Spatial analysis was conducted using the MuSpAn package (v1.1.1) [50] using Python (v3.10). Code available: https://github.com/ewestlundicr/MuSpAn---HPC.

To identify recurrent cellular neighbourhoods (RCN) a neighbourhood_enrichment_matrix was generated using the ms.networks.cluster_nneighbourhoods function. This was implemented using the K nearest neighbour (KNN) network type, using the minibatchkmeans clustering method. The number of nearest neighbours was set to 10. Following a series of optimisation tests, the number of clusters to identify was set to 8.

RCN CD8 enrichment was defined as the percentage of CD8+ cells inside a given RCN minus the percentage of CD8+ cells outside the RCN. CD8 enrichment was considered for n=143 samples. For each RCN, a paired Wilcoxon test considered the percentage inside vs outside for each sample, p-values were adjusted for multiple testing with the Benjamini-Hochberg method. The median enrichment score for each RCN was calculated for plotting.

Samples included in both the spatial and transcriptomic analysis were studied to identify differences in RCN abundance in ESTIMATE Stromal low (N=54) and high (N=59) samples. Wilcoxon p-values were calculated and adjusted for multiple testing with the Benjamini-Hochberg method.

To quantify direct cell-cell contacts in a 20 μm radius, a proximity network was generated for each sample using the msnetworks.generate_network function, khops = 1. To assess the proportion of cell contacts compared to random chance based on cell type abundance the observed / expected (O/E) ratio was calculated as (observed interaction frequency) / (expected interaction frequency under random null), where the expected frequency is the abundance of the target cell type in a given sample. To be included in the analysis, samples required a minimum of 50 source cells, a minimum of 50 target cells, and at least 10 contacts between the source and a given target. Interactions between source and target cells were considered if they occurred in a minimum of 10 independent samples.

### Imaging and quantification of the extracellular matrix

Slides were stained with picrosirius red as per the manufacturer’s protocol (Abcam, ab24632). In brief, slides were baked at 50°C overnight followed by an hour incubation at 60°C. Slides were deparaffinised and rehydrated. Nuclei were stained with Weigert’s Haematoxylin, collagen fibres were detected with Picrosirius red solution for 1 hour at room temperature, followed by two brief washes in acidified water (0.5% acetic acid in H_2_0). Slides were dehydrated, submerged in xylene and a cover slip applied.

Picrosirius red-stained slides were scanned using the Hamamatsu NanoZoomer scanner, with brightfield settings at x20 magnification. In keeping with the multiplex image ROIs, annotations were created using ImageScope x64 software (Aperio v12.4.6.5003). To avoid quantification of collagen features in muscle bundles, muscle bundles were visually excluded from the annotations.

Whole-slide images were processed into 2000×2000 pixel tiles with the Napari tool “Computational Preprocessing of extracellular matrix (c-pmat)” (https://github.com/IntegratedPathologyUnit-ICR/c-pmat/tree/main). Tiles outside of the annotated ROI or with less than 80% tissue were automatically discarded. Tiles were manually checked to ensure they were free of artefacts, muscle bundles or had insufficient tissue. Tiles were colour deconvoluted to extract the collagen fibres from the background cytoplasmic and nuclear features in Image J (v2.16.0). Brightness and contrast values were adjusted to remove remnants of nuclear features.

The Image J plugin TWOMBLI (v1) was used to quantify features of the extracellular matrix. TWOMBLI parameters were set following the published guidance [77]. Tile-level data was batch corrected using the ComBat function from the SVA R package (v3.54.0).

### Statistical analysis

Multimodal comparisons were conducted for all 124 tumour samples with available multiplex imaging, fibre tracing, and transcriptomic data.

Comparisons of ESTIMATE Stromal low and high groups were performed using unpaired Wilcoxon rank-sum tests. Where multiple features were assessed, corrections for multiple testing were applied using the Benjamini-Hochberg (BH) method as indicated.

Comparisons involving more than two groups were tested first with the Kruskal-Wallis method, if significant, subsequent pairwise comparisons were tested with the Wilcoxon method and adjusted for multiple testing with the BH method.

### Single-nuclear RNA sequencing analysis

#### Sample inclusion

Single-nuclear RNA sequencing (snRNA-seq) data from human bladder tumours were obtained from the processed .h5ad object published by Gouin *et al.* (2021) GSE169379 [18]. As the publicly available dataset had already undergone preprocessing and quality control by the original authors, no additional filtering based on transcript counts, mitochondrial gene content, or doublet detection was performed in this study.

To ensure robust estimation of communication probabilities in CellChat and to mitigate stochastic noise from low-density clusters, we applied a strict inclusion threshold: patients contributing fewer than five fibroblasts were excluded (n=3 excluded from an initial n=25). The final analytical cohort comprised 22 patients.

snRNA-seq data were processed using Scanpy (v1.12). The relative proportions of fibroblast subtypes were calculated in pandas (v2.3.3) for each patient by generating a contingency table of patients by subtypes counts, normalised to the total fibroblast count per patient. For visualisation, patients were ranked in descending order of PDPN fibroblast frequency and visualised using stacked bar charts Matplotlib (v3.10.8).

#### Data normalisation and integration

Raw counts were normalised to a target sum of 10,000 per nucleus and log(x+1) transformed. Following the Gouin et al. workflow, the top 1,500 highly variable genes (HVGs) were identified, batch effects were accounted for during HVG identification using the batch_key parameter.

Data integration was performed using scVI-tools (single-cell variational inference) within a Scanpy-based workflow. The model was initialised using the raw integer counts of the 1,500 HVGs, with "batch" specified as the categorical covariate to minimise technical variation across different runs while preserving biological signal. The scVI h model was configured with 20 latent dimensions (n_latent=20) and a single hidden layer, using a Zero-Inflated Negative Binomial (ZINB) gene likelihood distribution to model the sparse count distribution characteristic of single-cell RNA-sequencing data. The model was trained for a maximum of 200 epochs with a learning rate of 0.001 and a KL-divergence warmup of one epoch. To prevent overfitting, early stopping was implemented. Finally, a neighbourhood graph was constructed using the derived scVI latent space (X_scVI), identifying the 50 nearest neighbours for each cell to facilitate downstream clustering and manifold visualisation. The integrated data underwent dimensionality reduction using the UMAP algorithm (v0.5.11). UMAP embedding was calculated with a minimum distance (min_dist) of 0.3 and a spread of 2.0. Success of integration was determined by projection of batch and patient information onto the UMAP. In addition, fibroblast subtypes were visualised alongside the marker gene expression values to confirm clear delineation of distinct fibroblast subtypes (Figure S2G).

#### Differential Expression Analysis

Differential gene expression analysis (DEA) was performed between the fibroblast subtypes using the sc.tl.rank_genes_groups function in Scanpy. Differentially-expressed genes were identified using the Wilcoxon rank-sum test (method=”wilcoxon”). The pts=True parameter was enabled to calculate the percentage of cells expressing each gene within each cluster for visualisation. The top five marker genes for each subtype, ranked according to the Wilcoxon test statistic, were visualised using sc.pl.rank_genes_groups_dotplot).

#### Gene set enrichment analysis

Gene Set Enrichment Analysis was performed on the full transcriptome of each subtype using the GSEApy package (v1.2.1). Genes were ranked according to their differential expression test statistics. A negligible deterministic offset was applied to resolve tied ranking statistics while preserving the original gene ordering.

Pre-ranked GSEA was performed against the MSigDB Hallmark (2020) database. The analysis was restricted to gene sets with a size between 5 and 1,000 genes and performed with 1000 permutations to determine significance. For visualisation, pathways with an FDR q-value < 0.5 were retained. For each fibroblast subtype and gene-set collection, up to 10 positively-enriched pathways were selected by ranking normalised enrichment scores (NES) in descending order, while up to 10 negatively-enriched pathways were selected by ranking NES in ascending order. The selected pathways were visualised as horizontal bar plots, with bar length and direction representing the NES and pathway-specific FDR q-values displayed alongside the bars.

#### CellChat

Ligand–receptor communication analysis was performed using CellChat (v1.6.1). Normalised gene-expression data and the CellChat human ligand–receptor interaction database was used to identify overexpressed signalling genes and ligand–receptor interactions. Communication probabilities were inferred using computeCommunProb, and interactions involving cell groups represented by fewer than 10 cells were excluded. Pathway-level communication probabilities, aggregated interaction networks and signalling centrality measures were subsequently calculated.

Outgoing ligand–receptor interactions were considered between FAP or PDPN fibroblast populations (senders) and all other cell populations (receivers). Interactions were classified according to the CellChat signalling categories; Secreted Signalling, ECM–Receptor, and Cell–Cell Contact, and the relative proportion of interactions within each category was visualised using proportional bar charts.

### *In vivo* experiments

Animal experiments were performed in accordance with UK regulations under project licence PP6496249 and approved for the ICR by the institutional animal ethics committee review board (AWERB). For BBN963 experiments, 1 x 10^6^ BBN963 cells were suspended in 100 µl (50 µl PBS and 50 µl Matrigel) and subcutaneously injected into the flanks of C57Bl/6J mice (either wild-type (WT) or genetically modified with Rag2KO). Tumours were measured three times weekly using callipers and tumour volume calculated using V = (W2 x L)/2. Tumours were harvested for histology when they reached license limits (figure 6B, S6C).

#### Murine irradiation and tumour measurement

Radiation was administered when tumours reached a volume of 100-150 mm^3^. Mice were anaesthetised using isoflurane via inhalation at 4 L/min (oxygen 1.5 L/min), then transferred to a mouse bed in the small animal radiation research platform (SARRP, Xstrahl) and light anaesthesia maintained at 2 L/min. A cone beam CT image of the mouse was taken, and treatment planning performed to target the x-ray beams to the tumour. X-ray radiation was administered through a 10×10 mm collimator via equal and opposite beams, totalling 8 Gy at a dose rate of 2.5 Gy/min (∼90 secs per beam). Mice recovered in pre-warmed cages at 37°C. Mice were sacrificed and tissue collected when tumours were approaching licence limits for end-point survival studies (<18 mm diameter). Tumours were harvested from the flanks, bisected and fixed in 10% neutral buffered formalin for 24 hours. Fixed tumours were processed and embedded into paraffin blocks.

#### Murine immunohistochemistry and multiplex staining

Tumour sections (4 µm) were stained by IHC with CD8 antibody (eBioscience 14-0808 (4SM15)) performed on the Dako Autostainer Link48. Stained slides were scanned on a Zeiss Axio Scan.Z1 in brightfield using a 20x objective.

Multiplex staining of mouse tumours was performed as described above for human samples. Antibody information for the mouse CAF PhenoImager multiplex panel is provided in Table S5.

### Graphics

Schematics were generated with Biorender. Bar and dot plots were generated using ggplot2 (v3.5.1). Plots for *in vivo* experiments were generated in GraphPad Prism (v10.5.0). Heatmaps were generated using the pheatmap R package (v1.0.12), the clustering was calculated using Euclidean distances and the Ward method.

## Acknowledgements

This work uses data provided by patients and collected by the NHS as part of their care and support. We would like to thank the patients, families and clinical trials teams that contributed to the tissues included in this research. We also gratefully acknowledge the Breast Cancer Now Histopathology core at The Institute of Cancer Research for their assistance with H&E and IHC staining of mouse BBN963 tumours.

## Contributions

AB and AW wrote and edited the manuscript. AB, AW, NJ and AM conceptualised the research project. AB conducted experimental work and analysis. KO conducted animal studies and associated data analysis. MD conducted bioinformatic analysis of snRNAseq data. CS supported PhenoCycler image generation. MCC conducted analysis of TME multiplex immunofluorescence and PhenoCycler data. BG conducted pathological review. AL supported transcriptomic analysis. EW supported MuSpAn spatial analysis. PN supported extracellular matrix analysis. MP and EA generated murine multiplex immunofluorescence images. MRD and EA supported the animal husbandry for the Rag2KO study. RF, TS, TL supported the generation of human multiplex immunofluorescence images. MZ, KC, RTB, AC, KJ, SH, RH, SH, EH, NJ designed and coordinated the clinical trials and provided access to clinical material and data. All authors reviewed and commented on the manuscript.

