## Supplementary Figures & Tables for "Spatially-resolved multimodal profiling identifies functionally-heterogeneous cancer-associated fibroblasts associated with poor radiotherapy outcomes in muscle-invasive bladder cancer"

### Supplementary figure 1:

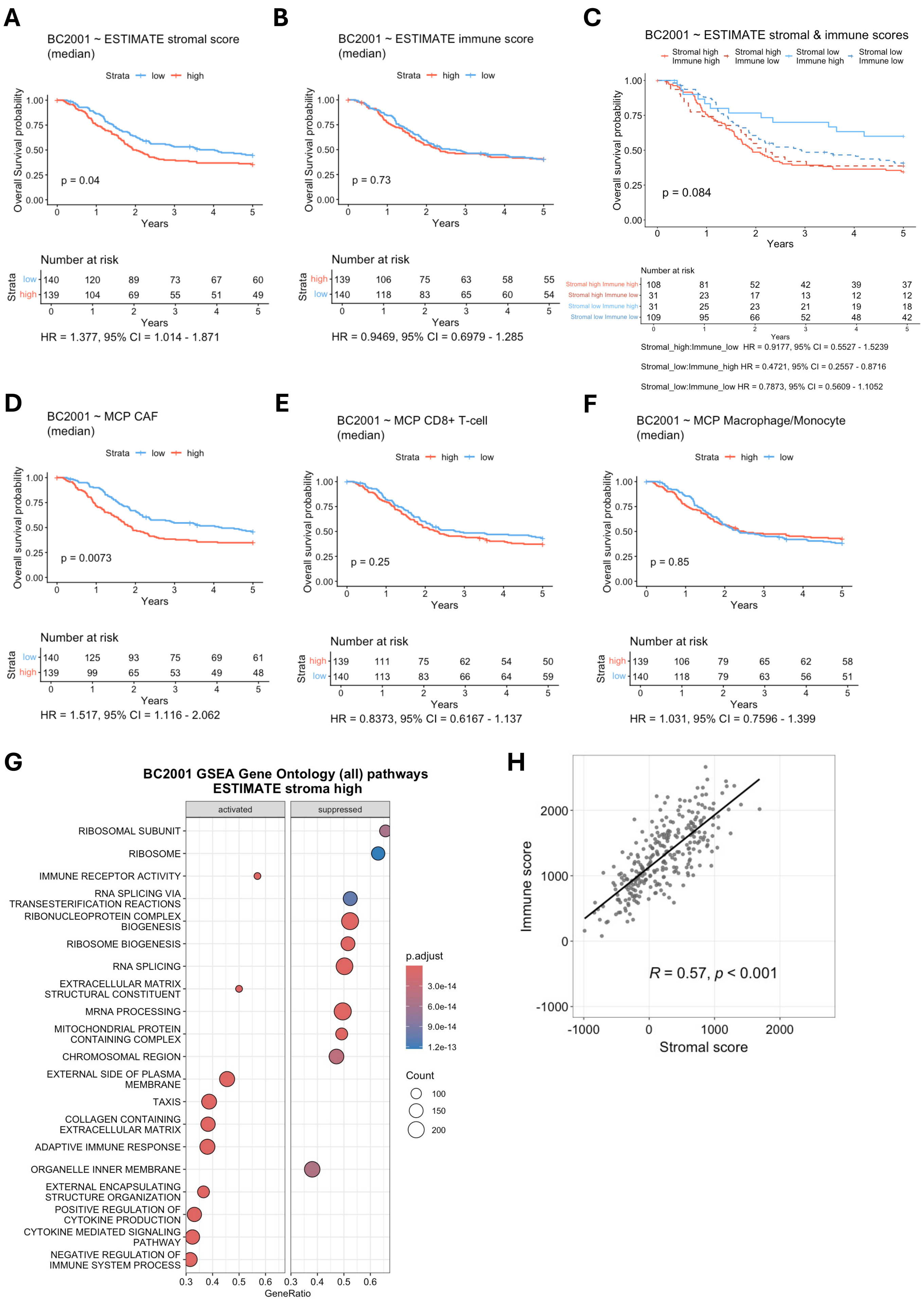

#### Supplementary Figures

##### Supplementary figure 1:

Kaplan-Meier curves of BC2001 overall survival dependent on the median cut point of: **A**. the ESTIMATE Stromal score, **B**. the ESTIMATE Immune score, **C**. a combination of the ESTIMATE Stromal and Immune scores, **D**. the Microenvironment Cell Populations (MCP)-counter CAF score, **E**. the MCP-counter CD8+ T-cell score and **F**. the MCP-counter Macrophage/Monocyte score. **G**. A dot plot of the top gene ontology pathways activated or suppressed in ESTIMATE Stromal high samples (optimal cut point). **H**. The correlation between the ESTIMATE Immune and Stromal scores.

### Supplementary figure 2:

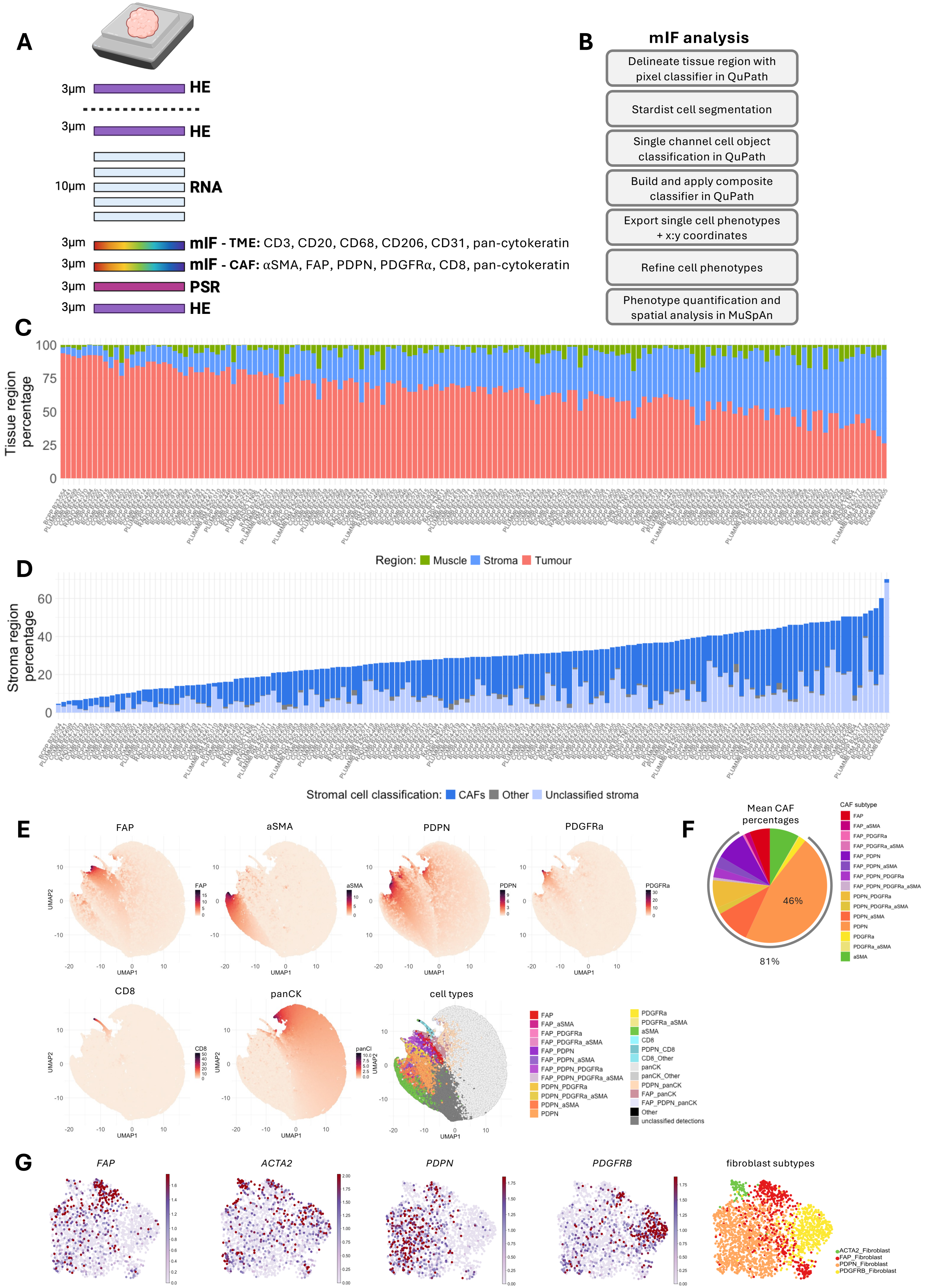

**Supplementary figure 2:**

**A**. A schematic of the tissue sectioning schema including an initial haematoxylin and eosin (H&E) section for pathological assessment, a second H&E for orientation, five 10 μm sections for nucleic acid extraction and transcriptomic profiling, two 3 μm sections for staining with the TME and CAF multiplex panel respectively, one 3 μm section for picrosirius red staining to be conducted on the serial section following the CAF multiplex panel and a final H&E to assess the tissue remaining in the block. **B**. The multiplex immunofluorescence image analysis workflow. **C**. The percentage of pixels classified as tumour, stroma or muscle per image. **D**. The percentage of stromal regions present in each sample, with the relative proportion of cells classified as CAFs, other or unclassified stromal cells. **E**. UMAP embedding of 100,000 randomly selected cells from images stained with the 6-plex CAF panel, cell are clustered by cell mean intensity with the cell mean intensity of each marker and the assigned phenotypes displayed. **F**. The mean proportion of each CAF population expressed as a percentage of the total CAF population. PDPN+ CAFs are the most abundant, where PDPN only expression accounts for 46% of CAFs and PDPN combinations represent 81% of all CAFs. **G**. UMAP embedding of single-nuclear RNA sequencing data, clustered by gene expression with the gene expression of *FAP*, *ACTA2*, *PDPN* and *PDGFRB* plus the assigned fibroblast phenotypes displayed.

### Supplementary figure 3:

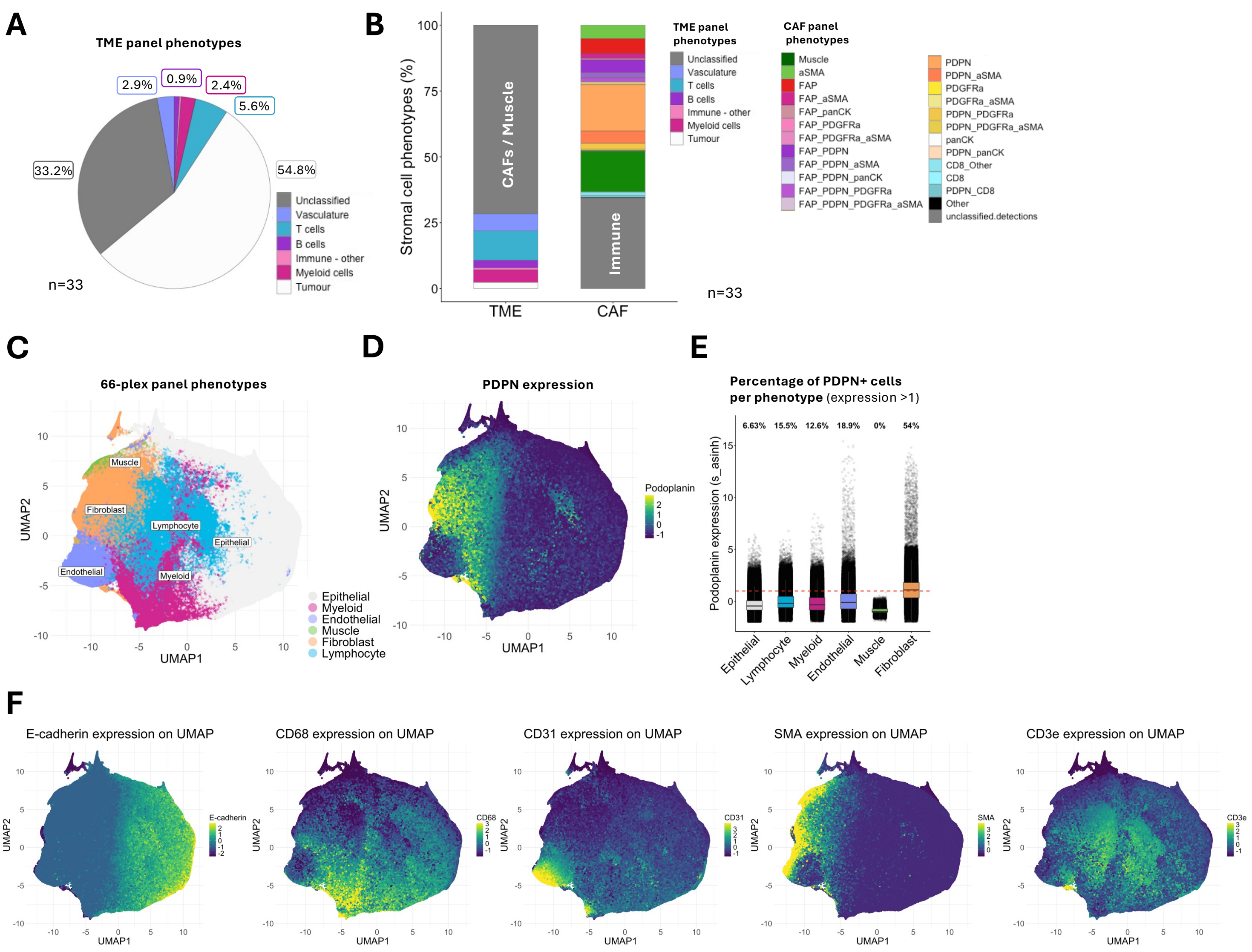

**Supplementary figure 3:**

**A**. A pie chart displaying the percentage of cell phenotypes detected by the TME multiplex panel. **B**. The proportion of cell phenotypes present in the stromal regions of images stained with the TME or CAF multiplex panel. To standardise comparisons, stromal regions in both panels are defined as pan-cytokeratin negative regions. The proportion of unclassified cells in the CAF panel likely represent the immune cells detected in the TME panel. **C**. A UMAP of all cells from images stained with the PhenoCycler 66-plex panel, cells are coloured by their assigned cell phenotype. **D**. A UMAP of all cells stained with the PhenoCycler 66-plex panel, cells are coloured by cell mean intensity expression of PDPN. **E**. The boxplot displays the PDPN expression detected in individual cells assigned to each cell phenotype. The dashed red line (y=1) indicates the threshold used to assign cells as PDPN+. Labels indicate the percentage of cells assigned to each phenotype that are above the threshold and are classified as PDPN+. **F**. UMAP embedding of all cells from images stained with the PhenoCycler 66-plex panel, cells are coloured by cell mean intensity expression of antigens used to assign define cell phenotypes.

### Supplementary figure 4:

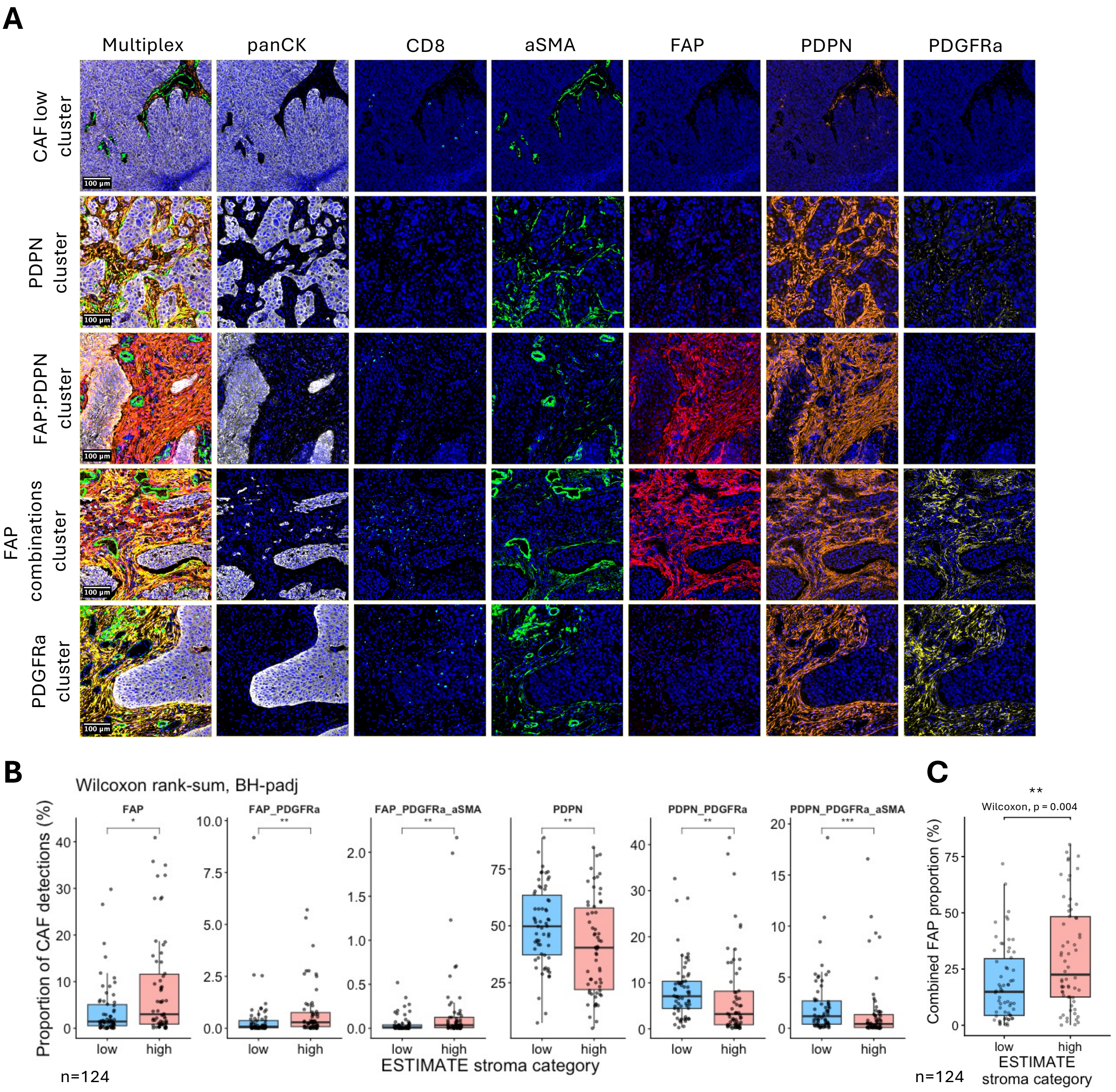

**Supplementary figure 4:**

**A**. Single channel immunofluorescent images of the examples shown in figure 3C. **B**. The proportion of individual CAF populations present in samples assigned as ESTIMATE Stromal low or high. Boxplots are shown for CAF populations that are significantly different. Benjamini-Hochberg adjusted Wilcoxon p symbols are displayed, $p<0.001$ ***, $p<0.01$ **, $p<0.05$ *. **C**. The proportion of combined FAP+ CAF populations present in samples assigned as ESTIMATE Stromal low or high, Wilcoxon $p=0.004$ **.

### Supplementary figure 5:

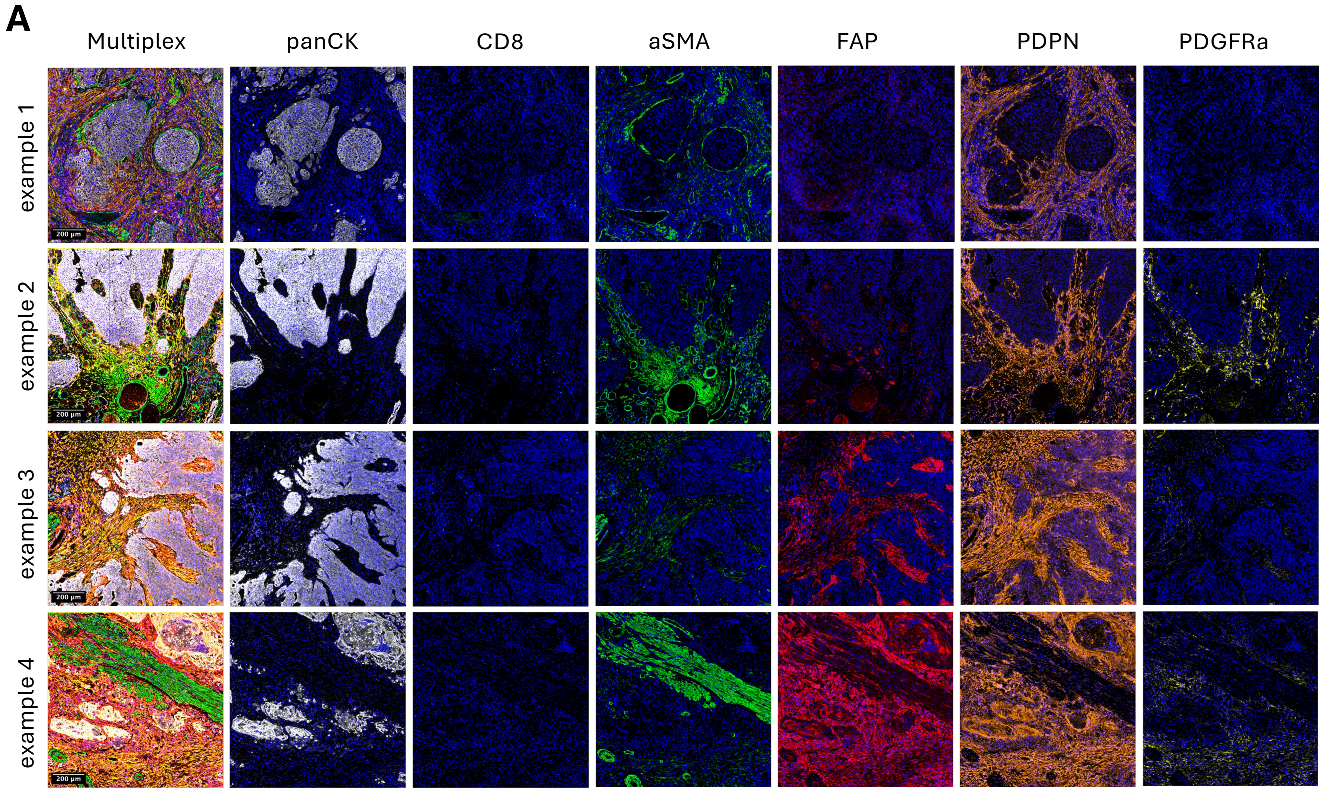

**Supplementary figure 5:**

**A**. Single channel immunofluorescent images of the examples shown in figure 4C.

#### Supplementary figure 6:

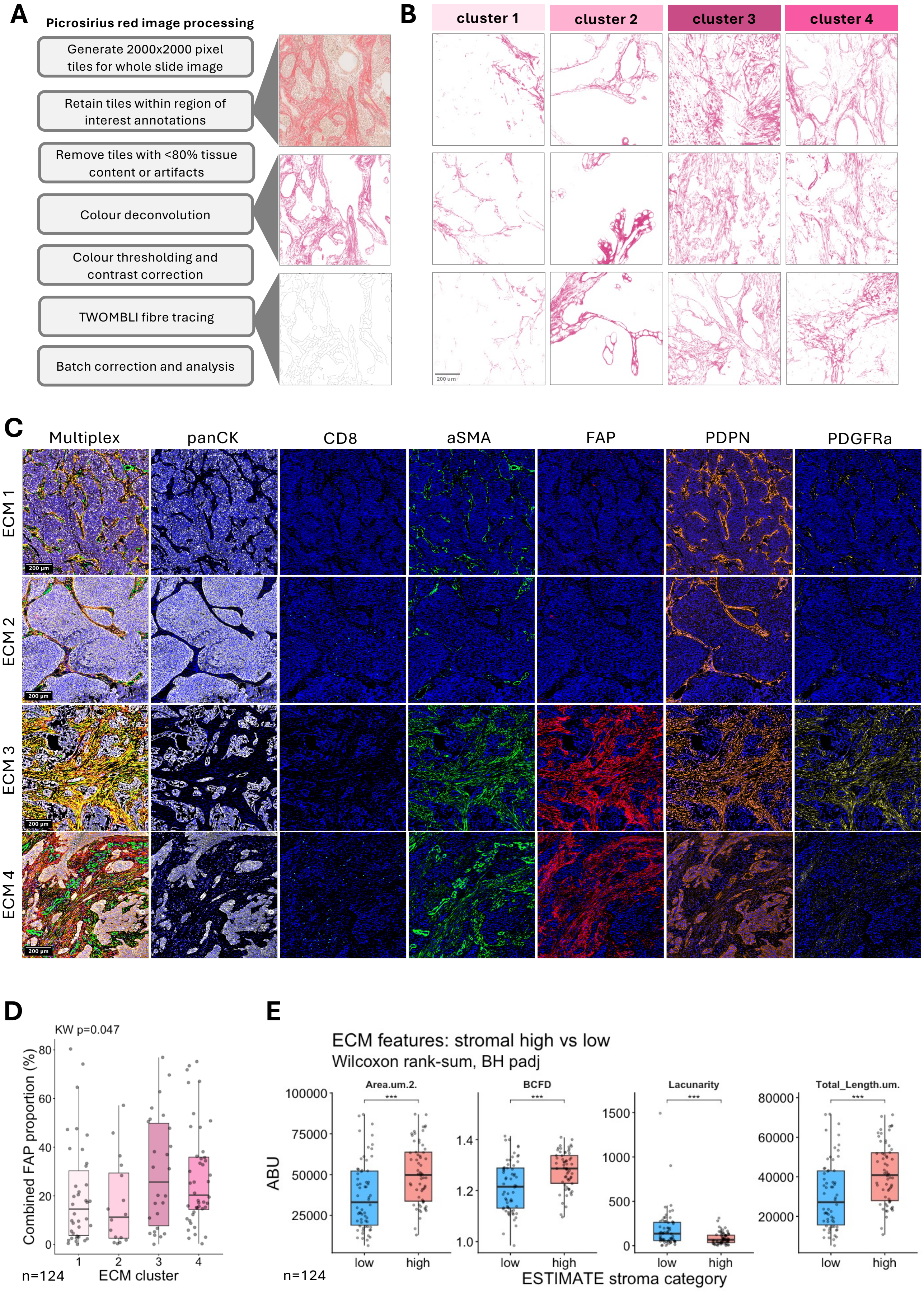

**Supplementary figure 6:**

**A**. A schematic of the processing steps to prepare picrosirius red stained images into tiles for TWOMBLI input. **B**. Additional representative images of tiles from samples assigned to each ECM cluster. **C**. Single channel immunofluorescent images of the examples shown in figure 5F **D**. A boxplot of the percentage of cells classified as CAFs in the multiplex images of samples assigned to each ECM cluster (n=124), Kruskal-Walis p=0.047. **E**. TWOMBLI features were compared between samples assigned as ESTIMATE Stromal low or high (n=124). Boxplots are shown for TWOMBLI features that are significantly different. Benjamini-Hochberg adjusted Wilcoxon p symbols are displayed, p<0.001 ***, p<0.01 **, p<0.05 *.

### Supplementary Figure 7:

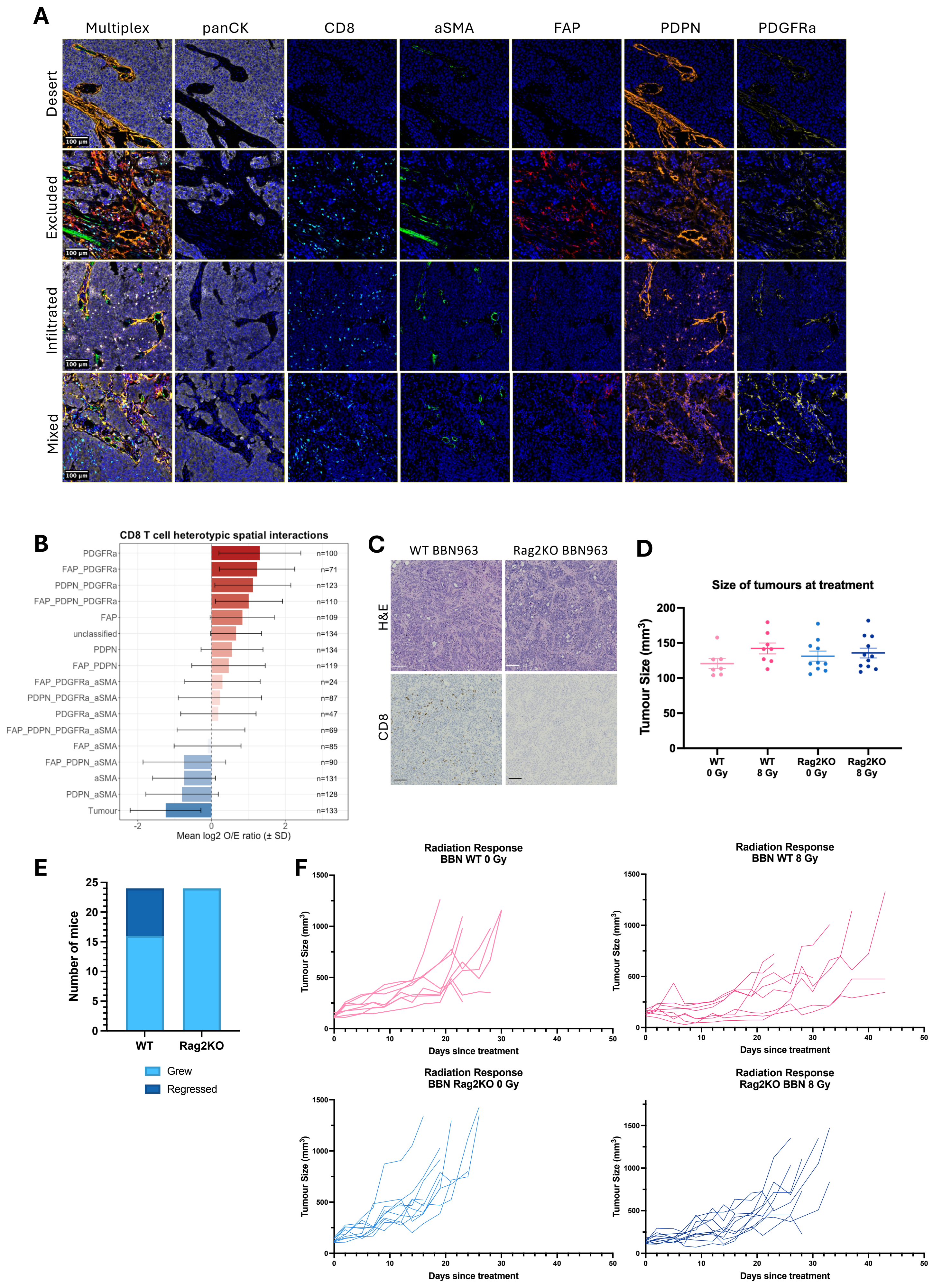

**Supplementary figure 7:**

**A.** Single channel immunofluorescent images of the examples shown in figure 6C. **B**. Observed / Expected odds ratio for heterotypic interactions between CD8+ cells (source) and CAF phenotypes or combined tumour cells (target). N indicates the number of samples the interaction could be assessed in. **C**. Representative images of untreated BBN963 tumours in wild-type (WT) or Rag2 knock out (Rag2KO) mice stained for haematoxylin and eosin (H&E) and CD8 by immunohistochemistry. **D**. Tumour measurements at start of treatment. **E**. Quantification of the number of BBN963 tumours that developed tumours or spontaneously regressed in WT or Rag2KO mice. **F**. Individual growth curves for WT and Rag2KO BBN963 tumours, untreated (0 Gy) and treated (8 Gy). Licence dictated end-point based on tumour size. Data was combined from 2 independent studies.

**Supplementary Tables**

**Supplementary table 1:** Clinical characteristics of BC2001 patients included in figure 1.

| Characteristic | Radiotherapy alone (n=157) | Chemoradiotherapy (n=122) | All patients (n=279) |
|---|---|---|---|
| **Sex** | | | |
| Male | 128 (81.5%) | 98 (80.3%) | 226 (81.0%) |
| Female | 29 (18.4%) | 24 (17.7%) | 53 (19.0%) |
| **Age - years** | | | |
| Mean | 71.6 | 70.4 | 71.1 |
| Range | 49-86 | 40-86 | 40-86 |
| **Grade** | | | |
| 1 | 0 | 1 (0.8%) | 1 (0.4%) |
| 2 | 25 (15.9%) | 15 (12.3%) | 40 (14.3%) |
| 3 | 130 (82.2%) | 106 (86.9%) | 236 (84.6%) |
| Unknown | 2 (1.3%) | 0 | 2 (0.7%) |
| **Pathological stage** | | | |
| 2 | 137 (87.2%) | 105 (86.1%) | 242 (86.7%) |
| 3a | 7 (4.5%) | 4 (3.3%) | 11 (3.9%) |
| 3b | 8 (5.1%) | 7 (5.7%) | 15 (5.4%) |
| 4a | 5 (3.2%) | 6 (4.9%) | 11 (3.9%) |
| **Radiotherapy schedule (Gy/fraction)** | | | |
| 55/20 | 46 (29.3%) | 40 (32.8%) | 86 (30.8%) |
| 64/32 | 111 (70.7%) | 82 (67.2%) | 193 (69.2%) |
| **Neoadjuvant chemotherapy** | | | |
| Yes | 38 (24.2%) | 36 (29.5%) | 74 (26.5%) |
| No | 119 (75.8%) | 86 (70.5%) | 205 (73.5%) |
| **Salvage cystectomy** | | | |
| Yes | 21 (13.4%) | 11 (9.0%) | 32 (11.5%) |
| No | 136 (86.6%) | 111 (91.0%) | 247 (88.5%) |

**Supplementary table 2:** Antibody information for the human PhenoImager CAF panel

| Antigen | Dilution | Antigen Retrieval | Clone, Supplier | Opal |
|---|---|---|---|---|
| FAP | 1:200 | pH9 | EPR20021, Abcam | 690 |
| CD8 | 1:500 | pH9 | C8/144B, Dako | 480 |
| PDGFRα | 1:200 | pH9 | EPR22059-270, Abcam | 570 |
| PDPN | 1:800 | pH6 | D2-40, Biolegend | 620 |
| αSMA | 1:5000 | pH6 | 1A4, Abcam | 520 |
| Pan-cytokeratin | 1:200 | pH6 | AE1/AE3, Dako | TSA/780 |

**Supplementary table 3:** Antibody information for the human PhenoImager TME panel

| Antigen | Dilution | Antigen Retrieval | Clone, Supplier | Opal |
|---|---|---|---|---|
| CD31 | 1:300 | pH9 | JC70A, Dako | 690 |
| CD68 | 1:150 | pH9 | PG-M1, Dako | 520 |
| CD3 | 1:100 | pH6 | SP7, Abcam | 570 |
| CD20 | 1:250 | pH6 | L26, Abcam | 480 |
| CD206 | 1:300 | pH6 | E2L9N, Cell Signaling Technology | 620 |
| Pan-cytokeratin | 1:200 | pH6 | AE1/AE3, Dako | TSA/780 |

**Supplementary table 4:** Antibody information for custom-conjugated antibodies added to the PhenoCycler Fusion IO60 panel.

| Antigen | Dilution | Clone, Supplier | Barcode | Fluorophore |
|---|---|---|---|---|
| FAP | 1:100 | EPR20021, Abcam | BX050 | AF647 |
| PDGFRα | 1:100 | EPR22059-270, Abcam | BX072 | AF647 |
| Collagen 1 | 1:500 | EPR7785, Abcam | BX078 | AF750 |
| Fibronectin | 1:200 | E5H6X, Cell Signaling Technology | BX106 | AF750 |
| Tenascin C | 1:100 | 4C8MS, Novus Bio | BX103 | AF750 |
| CD86 | 1:100 | E2G8P, Cell Signaling Technology | BX076 | AF750 |

**Supplementary table 5:** Antibody information for the mouse PhenoImager CAF panel

| Antigen | Dilution | Antigen Retrieval | Clone, Supplier | Opal |
|---|---|---|---|---|
| CD8a | 1:100 | pH9 | 4SM15, eBioscience | 570 |
| PDGFRα | 1:100 | pH9 | EPR5480, Abcam | 690 |
| FAP | 1:100 | pH9 | EPR20021, Abcam | 520 |
| αSMA | 1:2,500 | pH9 | EPR5368, Abcam | 620 |
| Cytokeratin 14 | 1:500 | pH9 | Poly19053, BioLegend | TSA/780 |
| PDPN | 1:200 | pH6 | 811, Antibodies online | 480 |

#### Supplementary Files

**Supplementary file 1:** QuPath script to stitch PhenoImager multiplex images post spectral unmixing.

**Supplementary file 2:** QuPath pixel classifier trained to detect regions of tumour, stroma and muscle in PhenoImager multiplex images stained with the Human CAF panel.

**Supplementary file 3:** QuPath composite object classifier trained to identify phenotypes of cells based on antigen expression in PhenoImager multiplex images stained with the Human CAF panel.

**Supplementary file 4:** The phenotype refinement key containing original phenotypes assigned by the QuPath composite classifier (supplementary file 3) and the adopted phenotype name for subsequent analysis.
