## Supplementary file 1 for "Spatially-resolved multimodal profiling identifies functionally-heterogeneous cancer-associated fibroblasts associated with poor radiotherapy outcomes in muscle-invasive bladder cancer": Stitch.groovy

```
//Modified by Colt Egelston and Jian Ye to do a directory of folders. 
//Change Line 34 to reflect folder locations. 

/**
 * Convert TIFF fields of view to a pyramidal OME-TIFF.
 *
 * Locations are parsed from the baseline TIFF tags, therefore these need to be set.
 *
 * One application of this script is to combine spectrally-unmixed images.
 * Be sure to read the script and see where default settings could be changed, e.g.
 *   - Prompting the user to select files (or using the one currently open the viewer)
 *   - Using lossy or lossless compression
 *
 * @author Pete Bankhead
 */

import qupath.lib.common.GeneralTools
import qupath.lib.images.servers.ImageServerProvider
import qupath.lib.images.servers.ImageServers
import qupath.lib.images.servers.SparseImageServer
import qupath.lib.images.writers.ome.OMEPyramidWriter
import qupath.lib.regions.ImageRegion
import qupath.lib.gui.dialogs.Dialogs

import javax.imageio.ImageIO
import javax.imageio.plugins.tiff.BaselineTIFFTagSet
import javax.imageio.plugins.tiff.TIFFDirectory
import java.awt.image.BufferedImage

import static qupath.lib.gui.scripting.QPEx.*

import static groovy.io.FileType.FILES

//*Change the below Folder Location!*
//def rootdir = new File("C:\\ComponentTIFF\\")
//alternatively could use 

def rootdir2 = Dialogs.showMessageDialog("Please navigate to componentTIFF directory","Please navigate to componentTIFF directory")
def rootdir = Dialogs.promptForDirectory(null)

//below is turend off for the purposes of batching folders in this script//
//Dialogs.promptForDirectory(//

def subdir =[]
rootdir.traverse(maxDepth:0){subdir.add(it)}

for (i in subdir){
    String s = i
    def dir = new File(s)
    print "Processing slide in folder " + dir
    def qupath = getQuPath()
    def files = []
    dir.traverse(type:FILES, maxDepth:0){files.add(it)}

    File fileOutput
    filename = s.split("\\\\")[-1]
    fileOutput = new File(rootdir,filename+'_Stitched.ome.tif')
    print "Output stitched file will be " + fileOutput
    
// Parse image regions & create a sparse server
print 'Parsing regions from ' + files.size() + ' files...'
def builder = new SparseImageServer.Builder()
files.parallelStream().forEach { f ->
    def region = parseRegion(f)
    if (region == null) {
        print 'WARN: Could not parse region for ' + f
        return
    }
    def serverBuilder = ImageServerProvider.getPreferredUriImageSupport(BufferedImage.class, f.toURI().toString()).getBuilders().get(0)
    builder.jsonRegion(region, 1.0, serverBuilder)
}
print 'Building server...'
def server = builder.build()
server = ImageServers.pyramidalize(server)

long startTime = System.currentTimeMillis()
String pathOutput = fileOutput.getAbsolutePath()
new OMEPyramidWriter.Builder(server)
    .downsamples(server.getPreferredDownsamples()) // Use pyramid levels calculated in the ImageServers.pyramidalize(server) method
    .tileSize(512)      // Requested tile size
    .channelsInterleaved()      // Because SparseImageServer returns all channels in a BufferedImage, it's more efficient to write them interleaved
    .parallelize()              // Attempt to parallelize requesting tiles (need to write sequentially)
    .losslessCompression()      // Use lossless compression (often best for fluorescence, by lossy compression may be ok for brightfield)
    .build()
    .writePyramid(pathOutput)
long endTime = System.currentTimeMillis()
print('Image written to ' + pathOutput + ' in ' + GeneralTools.formatNumber((endTime - startTime)/1000.0, 1) + ' s')
server.close()
}     


static ImageRegion parseRegion(File file, int z = 0, int t = 0) {
    if (checkTIFF(file)) {
        try {
            return parseRegionFromTIFF(file, z, t)
        } catch (Exception e) {
            print e.getLocalizedMessage()
        }
    }
}

/**
 * Check for TIFF 'magic number'.
 * @param file
 * @return
 */
static boolean checkTIFF(File file) {
    file.withInputStream {
        def bytes = it.readNBytes(4)
        short byteOrder = toShort(bytes[0], bytes[1])
        int val
        if (byteOrder == 0x4949) {
            // Little-endian
            val = toShort(bytes[3], bytes[2])
        } else if (byteOrder == 0x4d4d) {
            val = toShort(bytes[2], bytes[3])
        } else
            return false
        return val == 42 || val == 43
    }
}

/**
 * Combine two bytes to create a short, in the given order
 * @param b1
 * @param b2
 * @return
 */
static short toShort(byte b1, byte b2) {
    return (b1 << 8) + (b2 << 0)
}

/**
 * Parse an ImageRegion from a TIFF image, using the metadata.
 * @param file image file
 * @param z index of z plane
 * @param t index of timepoint
 * @return
 */
static ImageRegion parseRegionFromTIFF(File file, int z = 0, int t = 0) {
    int x, y, width, height
    file.withInputStream {
        def reader = ImageIO.getImageReadersByFormatName("TIFF").next()
        reader.setInput(ImageIO.createImageInputStream(it))
        def metadata = reader.getImageMetadata(0)
        def tiffDir = TIFFDirectory.createFromMetadata(metadata)

        double xRes = getRational(tiffDir, BaselineTIFFTagSet.TAG_X_RESOLUTION)
        double yRes = getRational(tiffDir, BaselineTIFFTagSet.TAG_Y_RESOLUTION)

        double xPos = getRational(tiffDir, BaselineTIFFTagSet.TAG_X_POSITION)
        double yPos = getRational(tiffDir, BaselineTIFFTagSet.TAG_Y_POSITION)

        width = tiffDir.getTIFFField(BaselineTIFFTagSet.TAG_IMAGE_WIDTH).getAsLong(0) as int
        height = tiffDir.getTIFFField(BaselineTIFFTagSet.TAG_IMAGE_LENGTH).getAsLong(0) as int

        x = Math.round(xRes * xPos) as int
        y = Math.round(yRes * yPos) as int
    }
    return ImageRegion.createInstance(x, y, width, height, z, t)
}

/**
 * Helper for parsing rational from TIFF metadata.
 * @param tiffDir
 * @param tag
 * @return
 */
static double getRational(TIFFDirectory tiffDir, int tag) {
    long[] rational = tiffDir.getTIFFField(tag).getAsRational(0);
    return rational[0] / (double)rational[1];
}
```
